# *Pyfiber*: an open source python library that facilitates the merge of operant behavior and fiber photometry- focus on intravenous self-administration

**DOI:** 10.1101/2022.09.02.506312

**Authors:** Dana Conlisk, Matias Ceau, Jean-François Fiancette, Nanci Winke, Elise Darmagnac, Cyril Herry, Véronique Deroche-Gamonet

## Abstract

**Background:** Advances in *in vivo* fluorescent imaging have exploded with the recent developments of genetically encoded calcium indicators (GECIs) and fluorescent biosensors. Their use with a bulk imaging technique such as fiber photometry (FP) can be highly beneficial in identifying neuronal signatures in behavioral neuroscience experiments.

Popularity of FP has grown rapidly. Initially applied to classical conditioning, its integration into operant behavior paradigms is progressing. However, in operant behavior, protocols can be complex including numerous scheduled events, while behavioral responses can occur in diverse and non-predictable manners. To optimize data processing and analysis, there is a need for a flexible tool to extract and relate behavioral and fiber photometry data occurring over operant sessions.

**New Method:** Applied to cocaine intravenous self-administration (using ImetronicⓇ polymodal apparati) and FP recordings in the prelimbic cortex (using Doric Lenses photometry system) in the rat, we established *Pyfiber*, an outline and open source data analysis python library that facilitates the merge of fiber photometry (using Doric Lenses) with operant behavior (using ImetronicⓇ). It allows relating activity changes within a neuronal population to the various behavioral responses and events occurring during operant behavior.

**Results:** We show some of the possibilities and benefits of the analytical tool *Pyfiber*, which helps to: 1. Extract the different types of events that occur in an operant session, 2. Extract and process the fiber photometry signals, 3. Select events of interest and align them to the corresponding fiber photometry signals, 4. Apply the most appropriate type of FP signal normalization and signal analysis according to the studied type of event or behavioral response, 5. Run data extraction and analysis on multiple individuals and sessions at the same time, 6. Collect results in an easily readable format for statistical analysis.

From our data and through the use of *Pyfiber*, we show that we can successfully record and easily analyze calcium transients surrounding events occurring during a cocaine self-administration paradigm in the rat.

**Comparison with Existing Method(s):** While other analytical tools can be used for streamlined fiber photometry analysis, they are either too rigid and specific or too flexible, requiring extensive coding to properly fit the data sets. Additionally, current tools do not permit easy exploration of multiple types of events in parallel- something that is possible with *Pyfiber*.

**Conclusions:** This work established an open source resource that facilitates the pairing of fiber photometry recordings (using Doric Lenses photometry system) with operant behavior (using ImetronicⓇ polymodal apparati), setting a solid foundation in analyzing the relationship between different dimensions of operant behavior with fluorescent signals from brain regions of interest.

## INTRODUCTION

There has been remarkable progress in the last twenty years regarding tools that can be used to monitor activity in specific neuronal populations using fluorescent indicators, especially applied to rodent models. The emergence and continuous improvement of genetically encoded calcium indicators (GECIs), specifically, GCaMPs, have allowed monitoring of specific neuronal populations in real time through fiber photometry (Nakai et al., 2001). These proteins are expressed in neurons and emit fluorescence when calcium binds (during the cytoplasmic calcium influx that occurs due to membrane depolarization opening voltage gated calcium channels).

In parallel, recent developments in genetically encoded biosensors permit further dissection of neural activity by observation of various markers (W. Wang et al., 2019). In addition to GECIs, genetically encoded voltage sensors (to observe membrane depolarization directly), genetically encoded pH sensors (to observe the pH changes due to the fusion of the synaptic vesicle with the plasma membrane during exocytosis/neurotransmitter release), and genetically encoded neurotransmitter indicators (to observe the neurotransmitter level in the synaptic cleft after vesicle release) have been developed and proven to work quite efficiently (Y. Wang et al., 2021). This, paired with expertise in viral genetics has permitted selective expression of these GECIs in subtypes of neuronal populations allowing researchers to observe alterations of specific neuronal subtype’s activity *in vivo*.

The use of these fluorescent indicators with a bulk imaging technique can be highly beneficial in identifying neuronal signatures in behavioral neuroscience experiments. Henceforth, the popularity for fiber photometry has grown rapidly- initially applied to classical conditioning paradigms (e.g., fear conditioning).

Its application to operant behavior is still limited. Technical constraints (eg. ensuring free expression of instrumental responses while head-connected) are now relatively minor difficulties, evidenced by the expansion of coupling of operant behavior with in vivo real time optogenetics: another optical fiber-based technique that is used to manipulate neuronal activity. However, for neural imaging, a main challenge for behavioral neuroscientists is to optimize data analysis by dealing with analytical constraints. In operant behavior, protocols can be complex and include numerous scheduled events. Behavioral responses can add to this complexity, with distribution occurring in a diverse and non-predictable manner. Due to this complexity, most behavioral neuroscientists limit themselves by focusing on analyzing one type of relatively well-controlled response or event.

Extraction of the most information from fiber photometry recordings during operant tasks relies on the ability to successfully apply the following steps:

1. Extract the different types of scheduled and unscheduled events that occur in the operant session
2. Extract and process the fiber photometry signals
3. Select all events or behavioral responses of a given type either globally or within a given time interval of the operant session
4. Align events of interest to corresponding fiber photometry signals
5. Apply the most appropriate type of signal normalization and signal analysis to study the neuronal response associated with a given type of event or behavioral response
6. Run data extraction and analysis on multiple sessions at same time
7. Collect results in an easily readable format for statistical analysis

In this objective, we created *Pyfiber*, a Python library that extracts and relates the data from our operant (ImetronicⓇ polymodal operant boxes) and fiber photometry systems (Doric Lenses). The library can be adapted to other behavioral and fiber photometry systems. *Pyfiber* can be integrated in a homemade command-line application and interfaced in a notebook. For this we recommend the Anaconda environment and Jupyter notebook (see details in the Methods).

By bridging the analytical gap between behavioral events occurring in an operant behavior paradigm with fiber photometry signals, the use of *Pyfiber* and its adaptation to other set-ups open the door to an exhaustive analysis of the neuronal signature of the events occurring while freely behaving animals are tested in complex protocols.

## MATERIAL AND METHODS

### Subjects

See Supplemental Online Materials (SOM)

### Surgeries

See SOM

### Histology

See SOM

### Intravenous self-administration training

See SOM

### Fiber photometry system

Fiber photometry recordings were done with a 1-site Doric Lenses photometry system (Doric Lenses, Quebec, Canada). Intensity of the 405 and 465 nm lights were sinusoidally modulated at 572 and 268 Hz respectively. Light was coupled to a filter cube (FMC4, Doric Lenses), converging onto a patch cord that was connected to the animal’s implanted optical fiber. Fluorescence collection was through the same patch cord, then passed to a photoreceiver (Visible Femtowatt Photoreceiver Module Model 2151, Newport). Output control and data acquisition was synchronized through a Doric Lenses Photometry Console, then passed to a PC that ran Doric Neuroscience Studio software. Data was low pass filtered with a cutoff frequency of 12 Hz.

### Intravenous self-administration (SA) apparatus

The self-administration setups were composed of plexiglas and metal (ImetronicⓇ, France). Each chamber (40 cm long × 30 cm width × 52 cm high) was encased within a larger opaque box equipped with exhaust fans that assured air circulation and masked background noise. During the sessions, animals were placed in a chamber where their chronically implanted intra-cardiac infusion catheter was connected to a pump-driven syringe. Two holes, located on opposite sides of the chamber at 5 cm from the grid floor, were used to record responding. The chambers were equipped with a blue cue light (1.8 cm in diameter) located on the opposite wall from the active hole at 33 cm from the grid, a white house light at the top of the chamber, and a white cue light (1.8 cm in diameter) located 9.5 cm above the active hole. Experimental contingencies were controlled and data collected with a PC-windows-compatible software (ImetronicⓇ, Marcheprime, France).

### Fiber recording during intravenous self-administration

The rats were connected to a dual pharmacology/optic fiber-motorized commutator (ImetronicⓇ, Pessac, France) which ensured that the patch cord and the line containing the cocaine solution rotated smoothly and in parallel to prevent tangling of the two lines **(Figure S1)**. The ImetronicⓇ behavioral system was connected to the Doric Lenses fiber photometry system by a TTL output. This TTL output initiated and stopped the fiber photometry recordings.

The behavioral protocol was identical to the basal self-administration training. The session consisted of three drug periods (40 min) separated by two no drug periods (15 min). Drug periods were characterized by illumination of the chamber by a blue light (LED2) while no drug periods were characterized by illumination of the chamber by a white house light (HLED) **(Figure 1A)**. Inactive nose-pokes during the duration of the sessions were recorded but had no scheduled consequences and active nose-pokes during the no drug period were recorded but had no scheduled consequences. An FR5 schedule with a maximum of 35 self-infusions was applied. Once the animals reached the 5 necessary nose-pokes in the active hole, the white cue light (LED1) illuminated for four seconds, with the pump being activated (delivering 46 microliters, 0.8 mg/kg/inf) one second after the illumination of the white light. Then, both the white cue light and blue light were turned off during the 40 second timeout period **(Figure 1B)**. Active and inactive responding during the timeout period was recorded but had no scheduled consequences.

**Figure 1:**
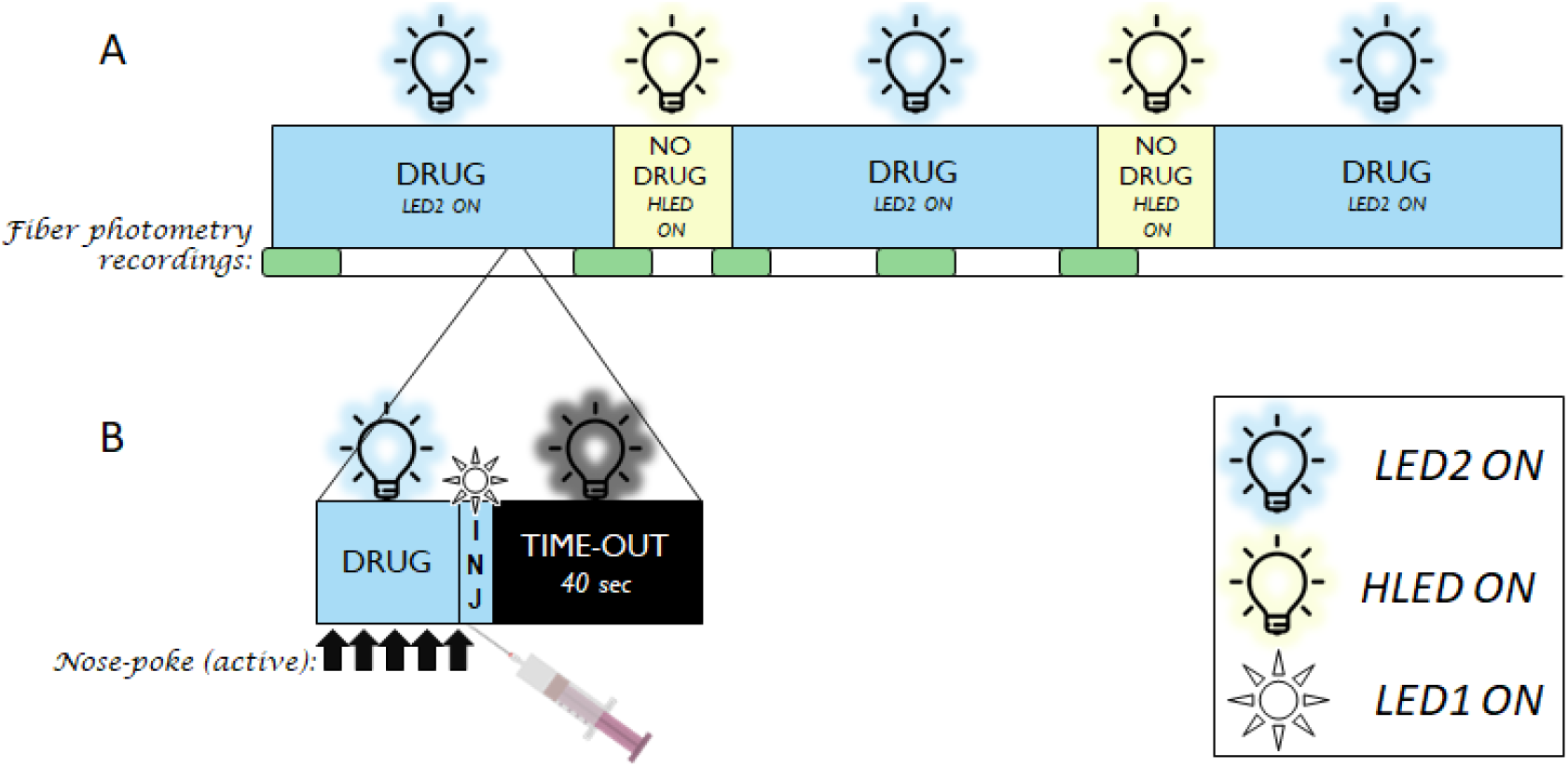
Schematic of self-administration protocol: A. The self-administration sessions are 2 hours and 30 minutes in entirety, consisting of three 40-minute drug periods (characterized by illumination of LED2) separated by two 15-minutes no drug periods (characterized by illumination of the HLED). The fiber photometry recordings are indicated by the green blocks shown underneath the self-administration schematic. B. The ratio of responding was FR5, meaning that the rat needs to nose-poke in the active hole 5 times to receive an injection. Once the fifth nose-poke is done, the conditioned stimulus (LED1) illuminates and is paired with pump activation and drug infusion. A time-out period of 40 seconds occurs thereafter where there are no lights illuminated.

Fiber recordings may take place at any moment throughout the operant protocol as they are initiated and stopped by the TTL output from the ImetronicⓇ system. The duration of the recordings in our experimental protocols ranged from 60 sec to 16 minutes depending on our experimental questioning. We give here examples for fiber recordings taking place at the initiation of the self-administration session in the first drug period (D1), around the shift from D1 to the first no drug period (ND1), around the end of ND1 to the second drug period (D2), in D2, and the shift from D2 to the second no drug period (ND2) (green boxes on **Figure 1**).

### Behavioral data processing (ImetronicⓇ)

Behavioral data is outputted by ImetronicⓇ as a tab-separated values text file, where each row represents a different message encoded by integers in each column. The first step consists of extracting and translating the data. The information retrieved are timestamps for a number of different events (nose-pokes in active and inactive holes, injections, injection associated conditioned cue light stimulus (CS), drug and no drug period-associated discriminative stimuli (DS) and house light switches).

From this raw data, specific events and intervals are computed. First, intervals corresponding to the different periods are algorithmically retrieved from the combination of light activation, and event /time-based rules. This strategy is particularly useful for intervals occurring at unpredictable times, such as timeouts following a drug injection, but also allows the accurate determination of timestamps for experimenter defined periods (such as the fixed drug and no drug periods), which is crucial for later alignment with fiber photometry data. Finally, a variety of more specific information (such as the position of each nose-poke in the fixed ratio series, or the first nose-poke in a given interval) are retrieved.

All of these events are defined and can be modified in the configuration file of *Pyfiber* (see Results- Configuration File).

Both the general extraction of events and intervals as well as custom experimenter-defined events and intervals exist due to the potential that some behavioral protocols have more complex events occurring.

For example, the general events and intervals are strictly defined by the data extracted by ImetronicⓇ (event = moment the house light turns on (hled_on), interval = the time period in which the house light is on (HLED_ON)).

In our protocol, the house light indicates the no drug period (**Figure 1A**). Normally, the switch from the drug to no drug period happens on the switch from the drug period (LED2 ON), however if the animal happens to be in a time-out period (**Figure 1B**), the switch from the drug to the no drug period will be from darkness to the no drug period (darkness to HLED ON), which should not be compared to recordings in which the switch is occurring from the drug period (LED2 ON to HLED ON). Due to this, we added custom criteria within the configuration file to specifically identify these two different types of light-switch occurrences.

All of these events are defined and can be modified in the configuration file of *Pyfiber* (see Results- Configuration File).

### Fiber photometry data processing (Doric Lenses)

#### Signal Processing

The exact methodology of normalization of the calcium-dependent and isosbestic channels differs between labs, therefore we implemented multiple ways of signal processing into our library.

##### ΔF/F

The current standard in the literature is based on a linear regression model (Bruno et al., 2021; Lerner et al., 2015). The assumption is that outside of the events that trigger depolarization of neurons and thus calcium release, the signal and control channels will be strongly correlated. Thus, in order to scale the control to the signal, a linear regression is performed on with data from the two channels with the least square method. The regression coefficients are then used to scale the isosbestic channel (control) to the calcium-dependent channel (signal) by using **Equation 1**. The resulting value corresponds to an activity ratio compared to baseline, although different factors such as random noise and calcium indicator decay may render it less straightforward.

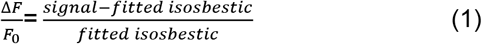

##### Z-scores

The second method consists of calculating Z-scores for the two signals, which can then be subtracted to obtain a unique normalized signal (**Equation 2**).

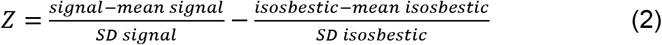

##### Peak analysis

In addition to the magnitude of the calcium-signal (reflecting a global increase in activation in the region of interest) the frequency of transients can also provide useful information. A transient is defined as a sudden increase in signal, which reflects neural activity.

However, due to random noise, artifacts and other constraints of fiber photometry, the normalized signal contains many local maxima, many of which are irrelevant. Thus, a strategy to determine if a transient is a peak, e.g. a significant transient reflecting an underlying synchronous depolarization. Many different methods exist in the literature. *Pyfiber* implements the most commonly used (Holly et al., 2019; Muir et al., 2018).

The signal is analyzed with a default of 10 second windows to limit the impact of signal decay (caused by photo-bleaching), which would lead to an underestimation of peaks towards the end of the recording. Standard scores are either calculated for the whole recording or inside each time window. Then, inside each window, the baseline median absolute deviation (MAD, or bMAD) is calculated and a first threshold is set (most commonly 2 to 2.5 times the MAD, default of *Pyfiber*=2.5). Signal data below this first threshold is removed and a new threshold is based on the MAD of the remaining data (pMAD, most commonly equal to 3.5 times this second MAD, default of *Pyfiber*=3.5). All signal data higher than this second threshold are considered as transients. Transients correspond to groups of consecutive data points (especially with high recording sampling rate; in our case around 1000 Hz). The final step is then selecting the timestamps for the highest point in each group of consecutive points, which then correspond to the peaks.

#### Perievent analysis

A major aim of the combination of operant behavior and fiber photometry is to determine whether the neural activity before or after a particular event (for example a light switch, a nose poke triggering a cocaine injection) is significantly different than at baseline, as well as comparing between subjects. In order to evaluate this, additional data processing surrounding events (perievent analysis) is performed.

##### Z-scores

To do so, Z-scores (**Equation 3**) are calculated by taking the time preceding the event as the baseline. Depending on which method was initially used for signal processing, perievent Z-scores can be generated from either the ΔF/F or Z-score.

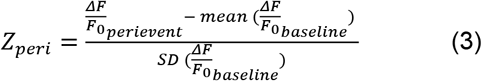

##### Robust Z-scores

For dispersed data, outliers can have a detrimental effect on both the mean and the standard deviation. Thus, another method has been proposed (Bruno et al., 2021). It utilizes robust Z-scores (**Equation 4**), which are similar to classical Z-scores but use the median instead of the mean and the median absolute deviation (**Equation 5**) instead of the standard deviation. This has the effect of removing the influence of outliers on the analysis.

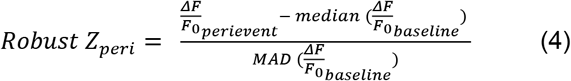

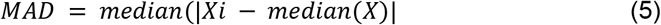

## RESULTS

### Pyfiber

#### Links

*PyPi*:

https://pypi.org/project/pyfiber/

*Read The Docs*:

https://pyfiber.readthedocs.io/en/latest/

*Gitlab*:

https://gitlab.com/inserm-u1215/pyfiber

#### 1. How to use *Pyfiber*

*Pyfiber* can be integrated in a homemade command-line application and interfaced in a notebook. For this we recommend the Anaconda environment.

##### 1.1. Installation of Anaconda, Jupyter Notebook, and *Pyfiber*

###### Anaconda and Jupyter Notebook

*Pyfiber* is recommended to be used through Jupyter Notebook. Guidance on how to install Anaconda and Jupyter notebook can be found here:

https://sparkbyexamples.com/python/install-anaconda-jupyter-notebook/

###### Installation of *Pyfiber*

Installation of *Pyfiber* can be done by opening the Anaconda Prompt and typing “pip install pyfiber”

##### 1.2. Importing *Pyfiber* to Jupyter Notebook

Before using any of the modules below, you need to import *Pyfiber* to the notebook. This can be done by calling the below command:

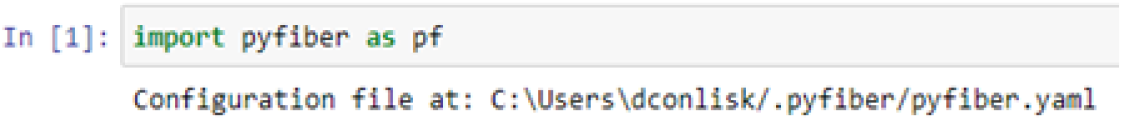

This command is done in all of the example notebooks that are provided on the Gitlab. When executed, it shows the location of the configuration file.

#### 2. Operating mode: Introduction to *Pyfiber* configuration file and operational modules

##### 2.1. Configuration file

The configuration file (in .yaml format (https://en.wikipedia.org/wiki/YAML) describes the different phases and events of a behavioral session (**see paragraph 3 below**): by modifying this file, it is possible to analyze any experiment that uses the Polymodal ImetronicⓇ system. It also describes the structure of the data from the fiber photometry system, as well as all the events of interest used in the analyses [e.g. shift from Drug to ND periods, nose-pokes (NP) during time-out (TO)…]. It also contains the default analysis settings, therefore changing the configuration file will alter default *Pyfiber* processes.

##### 2.2. The three operational modules

###### I. Analyze Module

The Analyze module is used to analyze the combination of ImetronicⓇ behavioral data and Doric Lenses fiber photometry data. It extracts identical behavioral responses or events occurring during fiber recordings, and performs fiber perievent analysis on them. However, it is not used independently- it is always called by *Pyfiber* when either the MultiSession or Session modules are called.

###### • MultiSession Module

The MultiSession module imports multiple sessions (each session as a folder containing one fiber data file (.csv) and its associated behavioral data file (.dat)) to perform group analysis. This permits simultaneous analysis of multiple recordings quickly and easily. The results can be exported to be graphed or for statistical analysis. This is the most complex function of *Pyfiber* and the most commonly used in our lab. For example, if the goal of the analysis is to see transients in response to a light presentation signaling cocaine delivery, MultiSession will be able to extract the time of every light presentation from the behavioral files and refer to each associated fiber photometry recording to perform perievent analysis.

###### • Session Module

The Session module imports a single session [one fiber data file (.csv) and one behavioral file (.dat)] for analysis of a single event. The Session module allows for perievent analysis of an indicated timestamp that can be extracted from the behavioral file. Therefore, if the interest is looking at a single event in a single session, the Session module is the most useful.

###### II. Fiber Module

The Fiber module is used by the analyze modules (both MultiSession and Session). This module is responsible for processing the fiber photometry data in preparation for perievent analysis.

However, if there is a specific interest in observing the fiber photometry data independently, the fiber module can be useful and can be used to analyze a single fiber photometry recording file (.csv).

###### III. Behavior Module

The Behavior module is also used by the Analyze module. It converts the raw ImetronicⓇ data file into files that compartmentalize the events and their times within the session in preparation for perievent analysis by the Analyze module(s). Like the Fiber module, the Behavior module can be used independently if there is a specific desire to look solely at behavioral events. It can be used in the two different following modes:

###### • Behavior Module

The Behavior module is used to look at a single raw ImetronicⓇ data file (.dat). This is used in both the MultiSession and Session modules to extract timestamps for the associated perievent analysis. If used independently, it can be used to extract behavioral responses or events from a single operant session.

###### • MultiBehavior Module

The MultiBehavior module is rather used to compare behaviors between subjects, groups or sessions using multiple ImetronicⓇ data files (.dat). It is used in the MultiSession module to visualize how the behavioral activities differ between the sessions that will be analyzed for perievent fiber photometry analysis.

#### 3. Description of the configuration file

Lines 1-9 of the configuration file contain general information regarding the identification tags and nomenclature used.

Line 4-5 indicates the automatic naming that was designed by our lab. It specifies that the folder containing the data files is formatted in a way that contains information about the experiment, rat number, experiment type, and self-administration session. For example, in the folder below, the experiment (AS21R), rat (12), experiment type (SA7), and session number (40) can automatically be extracted by *Pyfiber*.

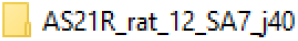

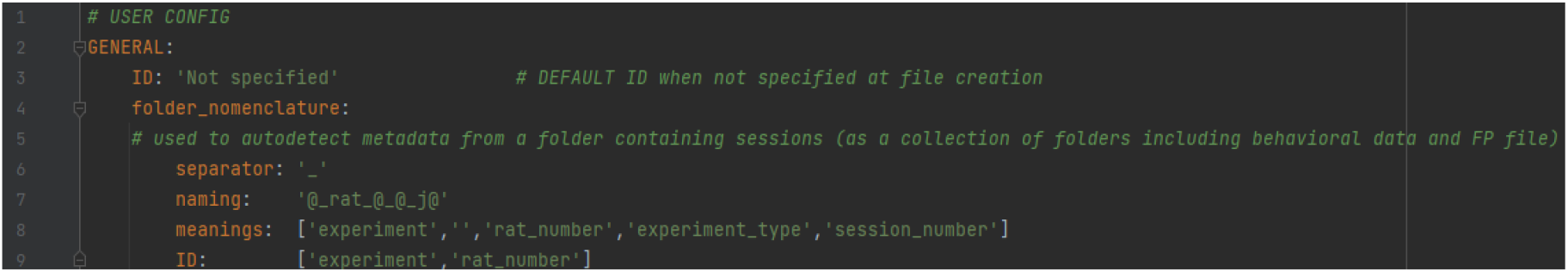

The default behavior file type (line 12) is ImetronicⓇ. As of now, there are no other possibilities. The default configuration of the behavior_time_ratio (line 13) is 1000, due to the behavior file containing information in milliseconds.

Lines 19-32 contain the definition of events that can be extracted from the raw ImetronicⓇ file and called for subsequent analysis.

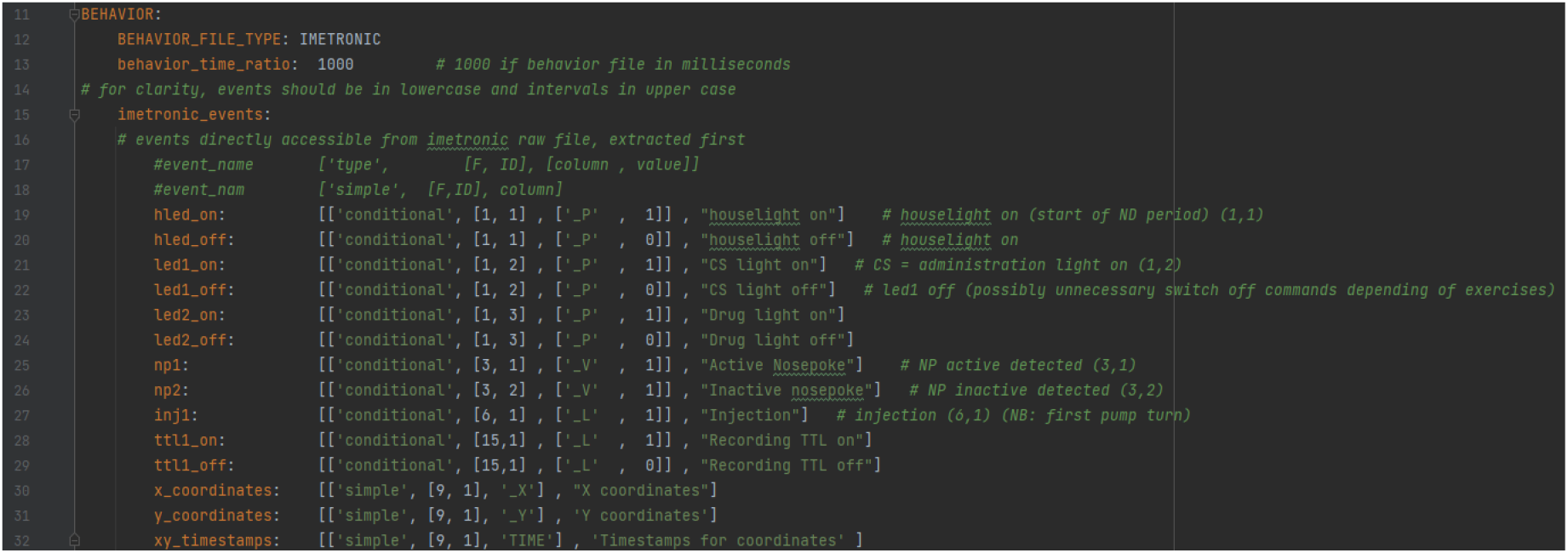

Along the same lines, there are intervals that are calculated from the aforementioned events that can be used as inclusion or exclusion criteria for analysis: for example, taking nose-pokes only when HLED is on.

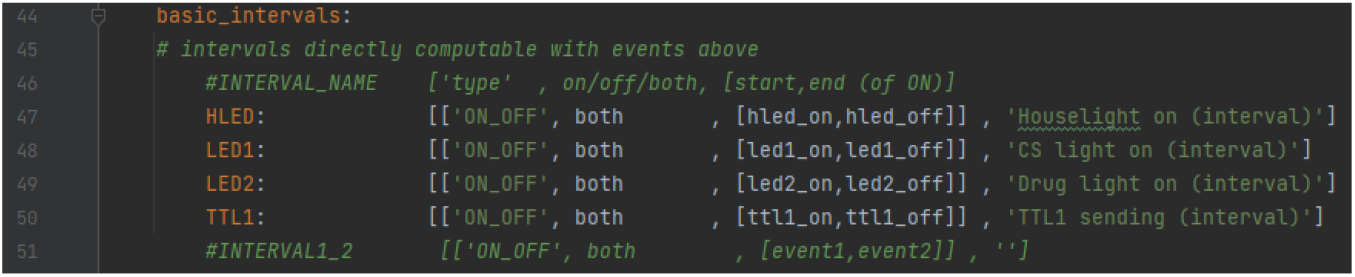

If there are other events or intervals of interest in analysis, such as levers or lickers, events can be added using the nomenclature found in rows 147-160. They just need to be added to the imetronic_events section.

For example, if levers are used instead of nose-pokes, there needs to be this addition to the imetronic_events section, that contains the imetronic configuration:

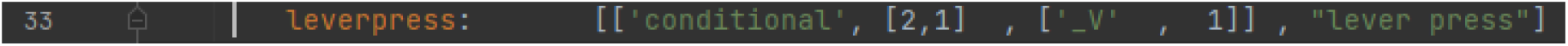

It is ‘conditional’ due to the fact that the determination of the timestamp of this event is conditional on a 1 occurring in the ‘_V’ column in the ImetronicⓇ file when the [F, ID] is equal to [2,1]. This information can be found in the ImetronicⓇ manual.

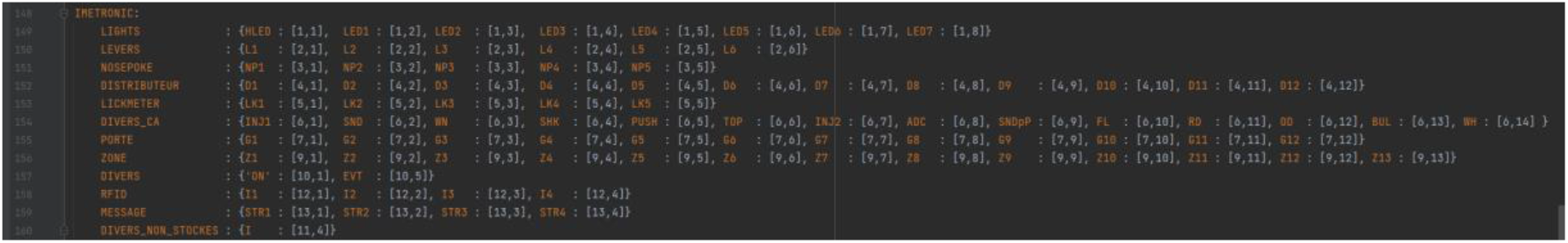

Additionally, within our protocol, there are complexities that require us to add additional custom events and intervals that can be found in lines 63-103. These include drug periods (LED2 ON), no drug periods (HLED ON), time-out periods (LED1 ON (cocaine conditioned stimulus) and the subsequent dark period) (**Figure 1A and B**).

Events that are present as TTL pulses in the fiber photometry data are planned to be able to be extracted, however it is not available in this current version.

In rows 106-126, there are the default plotting and color settings for events/intervals.

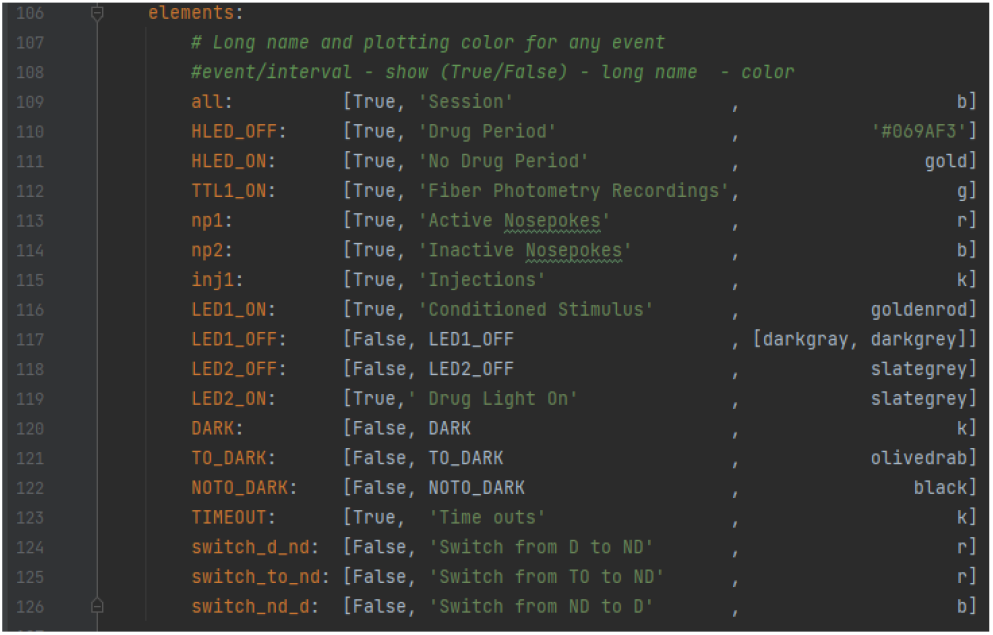

Next, the configuration file contains the default settings for the fiber photometry analyses. Changes in default settings can be done by changing this file and saving it. The next time you import *Pyfiber* to initialize analysis, the changed default settings will be taken into account. For example, if perievent analysis is always done on 5 seconds pre-event and 5 seconds post-event, change the white text in line 134 from [1.0, 1.0] to [5.0, 5.0].

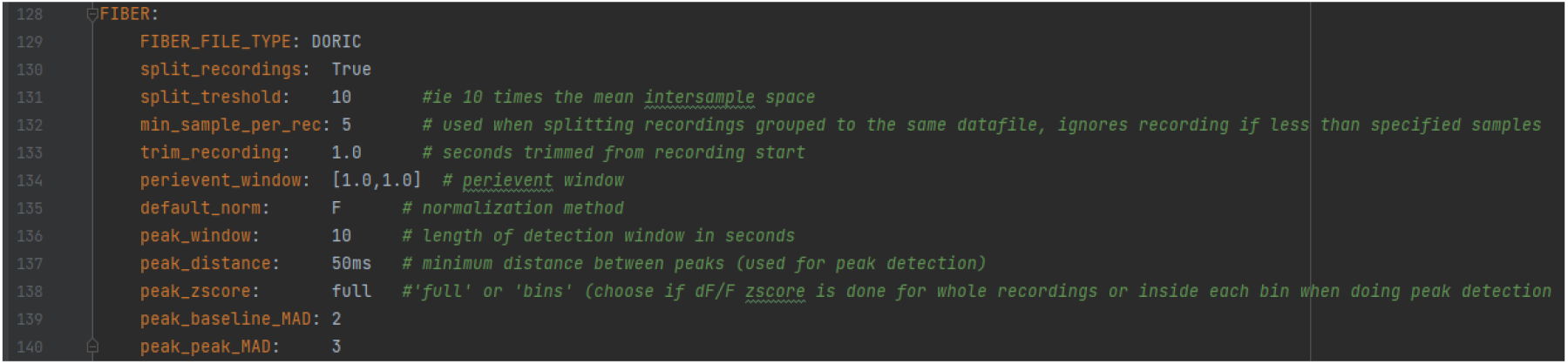

Lastly, the configuration file contains the nomenclature from the Doric Lenses system that is used to define the isosbestic, calcium dependent, time, and TTL channels, as well as other nomenclature that is used to define the columns containing time, signal, or TTL data. These can be found at the top columns of the .csv files generated by the Doric system.

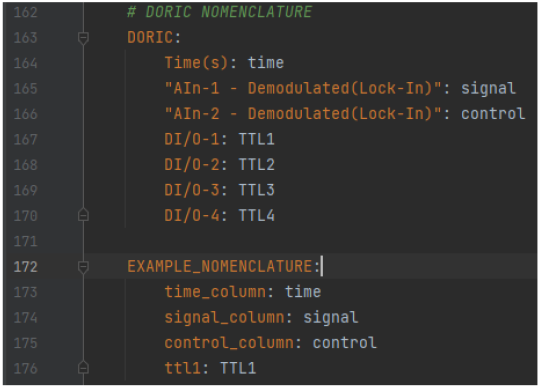

After confirming that the default settings are as desired, the modules can be called.

#### 4. Pyfiber application

For the sake of space, only tutorials on the Analyze module (MultiSession and Session) are shown here. For tutorials on Fiber and Behavior (MultiBehavior, Behavior) modules, please see the Jupyter Notebooks associated with those modules and the tables outlining the functions of the modules in the SOM.

##### 4.1 Multisession

As stated above, the MultiSession module is the prototypical use of *Pyfiber* when running fiber photometry recordings during an operant task over several sessions and/or on several individuals. It analyzes multiple behavioral and fiber data files in parallel. It uses the **analyze**, the **behavior** and the **fiber** modules.

A table can be found in the SOM that contains a selection of the most pertinent features and the commands associated with the MultiSession module.

Below is a tutorial explaining how we do our analysis with MultiSession. Two prototypical types of questionings are shown. The first analyzes a scheduled event (shift from no drug to drug period signaled by the shift between two distinct light conditions) while the second presents the analysis of non-scheduled events resulting from the animals’ behavior (in the present case: self-infusions).

###### 4.1.1. Calcium signals by the Prelimbic cortex principal neurons in response to the shift from the drug to the no drug period

The example is given on 3 recordings run during 3 independent behavioral sessions (all run on the same rat. The corresponding Jupyter Notebook can be found and has to be downloaded together with the csv (fiber data) and dat (behavior data) files at: https://gitlab.com/inserm-u1215/pyfiber/-/tree/main/docs/notebooks/MultiSession

The following steps shown are the only ones necessary for this example but are excerpts of a Jupyter Notebook containing a variety of analyses. The corresponding locations of the steps below in the MultiSession Jupyter Notebook are indicated in the comment just above the execution of code.

**Step 1***(MultiSession Jupyter Notebook: #1)*: Import *Pyfiber* in Jupyter Notebook.

**Step 2***(MultiSession Jupyter Notebook: #2)*: Creating a MultiSession object by calling pf.MultiSession() and indicating the file path in the parenthesis.

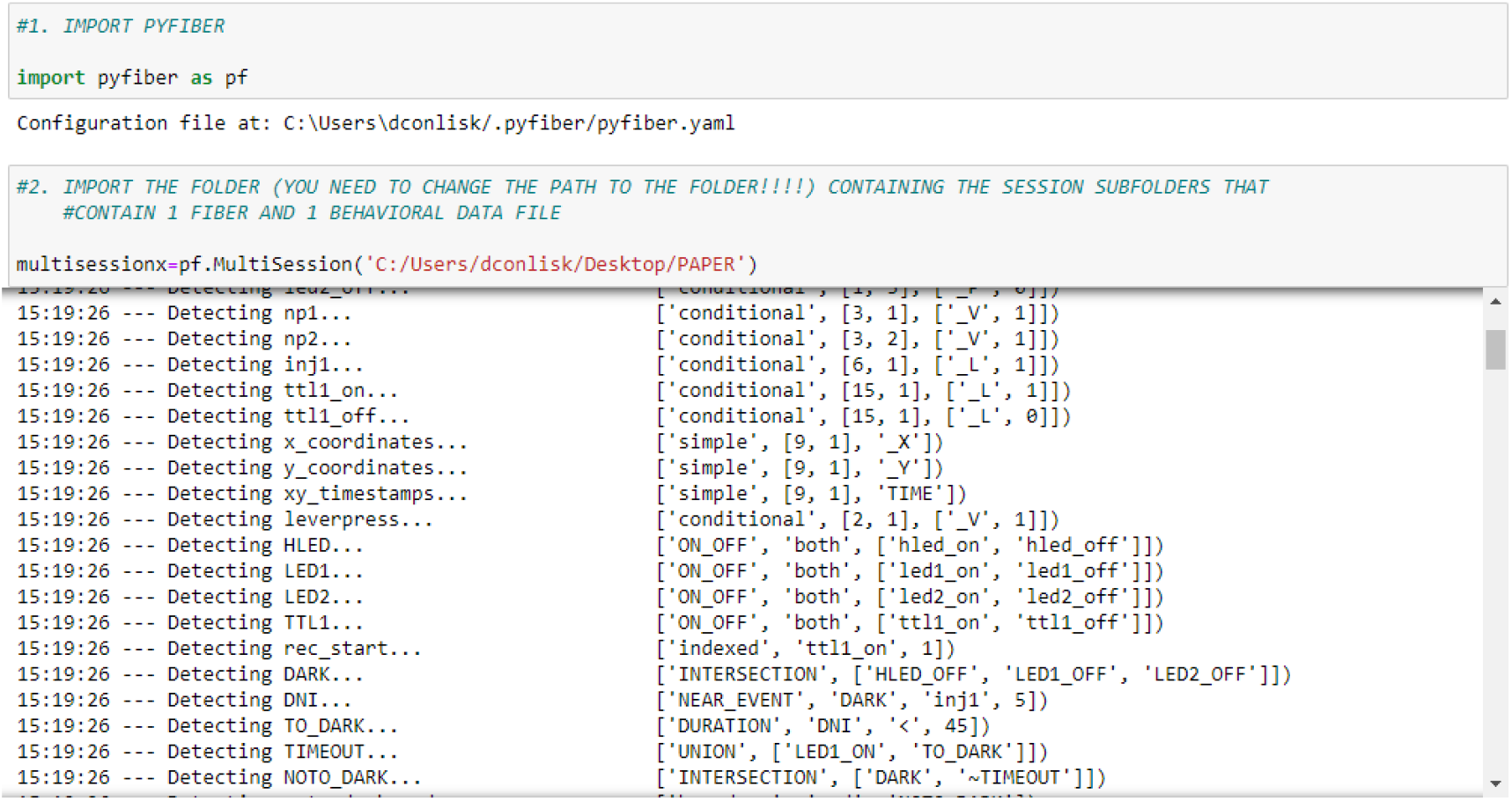

**Step 3***(MultiSession Jupyter Notebook: #8)*: Create a perievent object by calling .analyze from the MultiSession object. Indicate in the parenthesis which events should be used as the events of interest and the window that should be used surrounding the event.

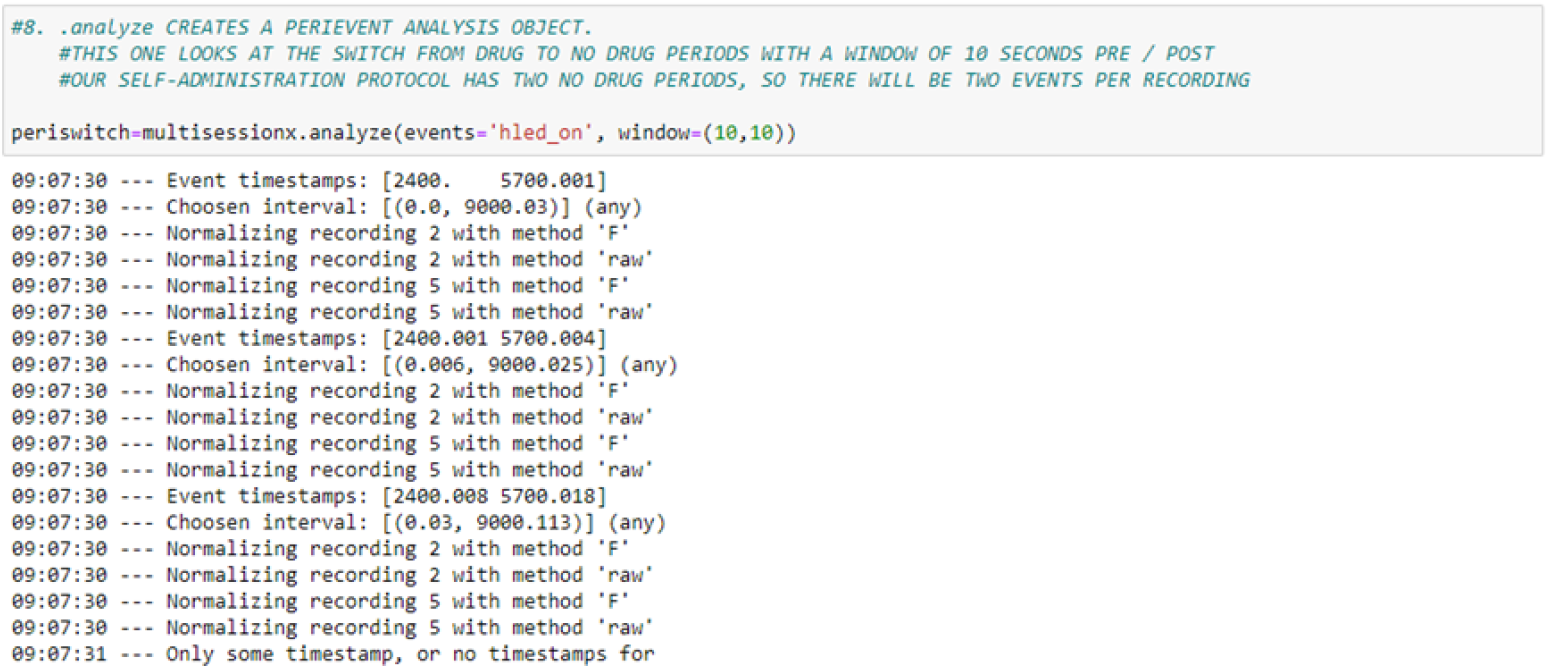

**Step 4***(MultiSession Jupyter Notebook: #9)*: Call .plot from the perievent object and indicate what to plot (shown here: processed signal) to see the individual (colored) and average (black) traces of the perievent signal.

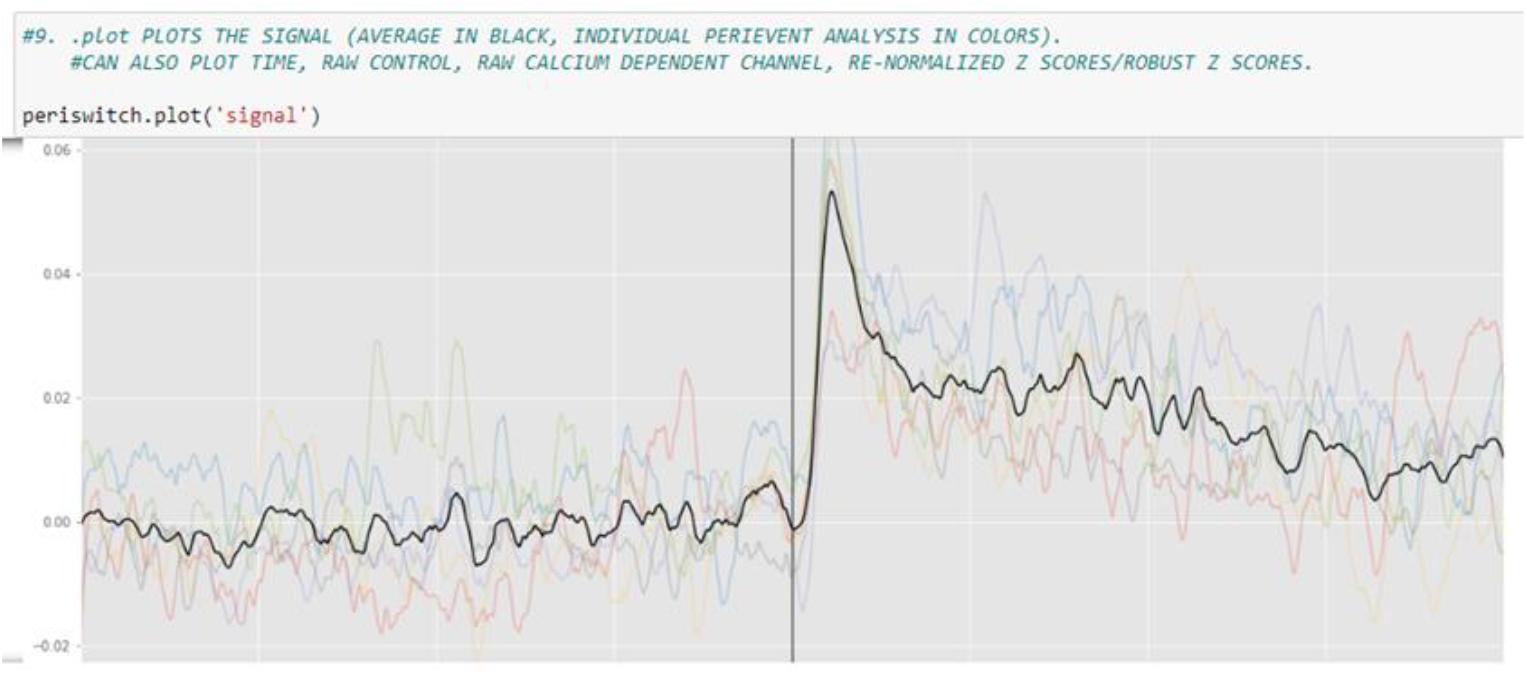

**Step 5***(MultiSession Jupyter Notebook: #10)*: Call .data from the perievent object to see a table containing the average values for the perievent signal: including pre/post AUC, dF/F, Z-Scores, Robust Z-Scores, peak frequency, peak amplitude, etc. For the full list of things that are in this table, see .data in the MultiSession table found in the supplementary material.

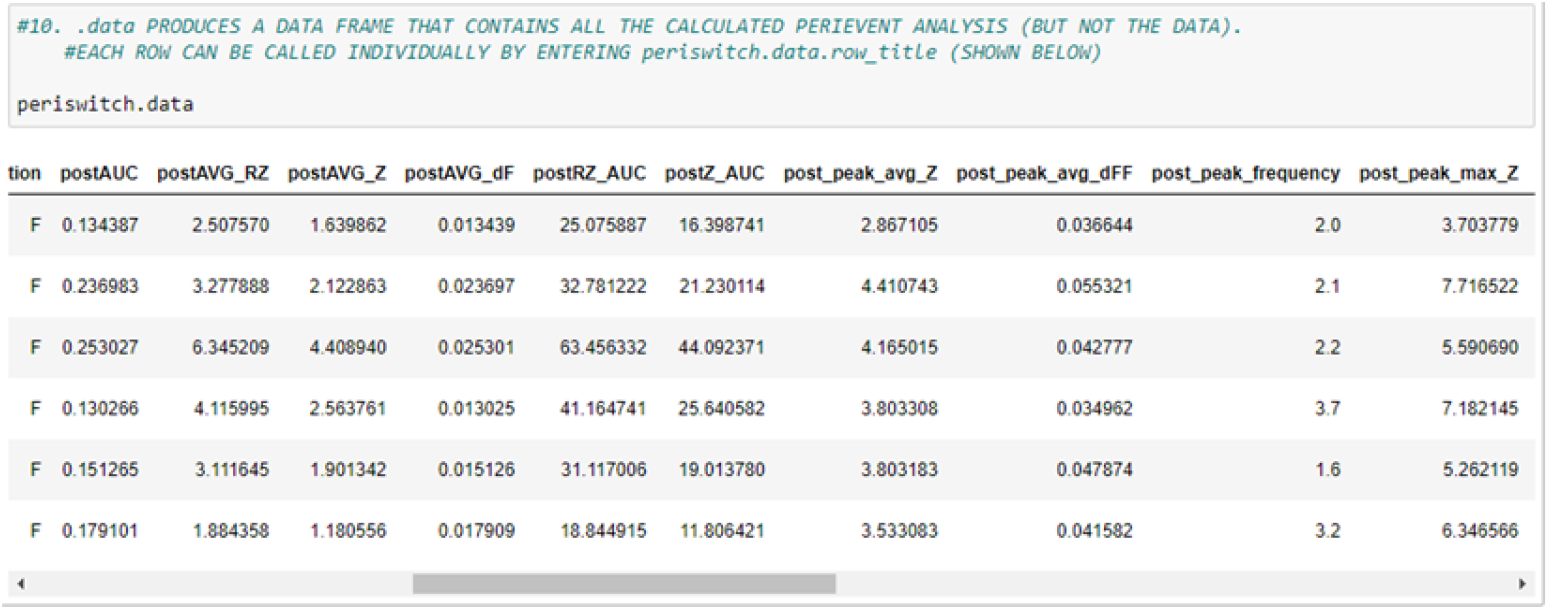

**Step 6***(MultiSession Jupyter Notebook: #12)*: Call .full_data from the perievent object to see a table that contains the dataframes for the perievent analysis.

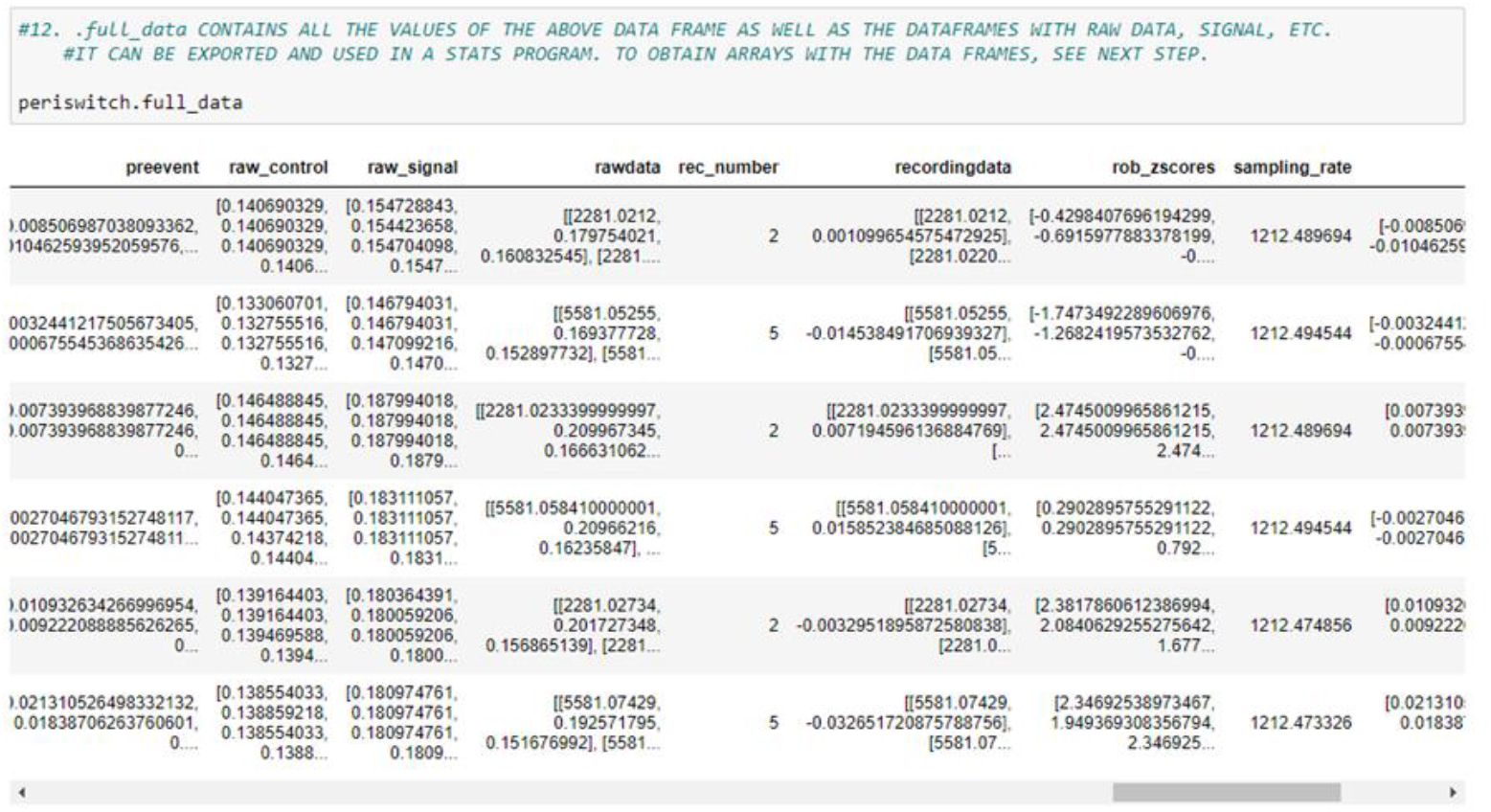

**Step 7***(MultiSession Jupyter Notebook: #15)*: Lastly, an example is shown of extraction of the dataframes containing the adjusted timestamps leading up to the event and the processed signal that corresponds to the timestamps. These can be exported to a .csv so the data can either be used in statistical programs or to create figures.

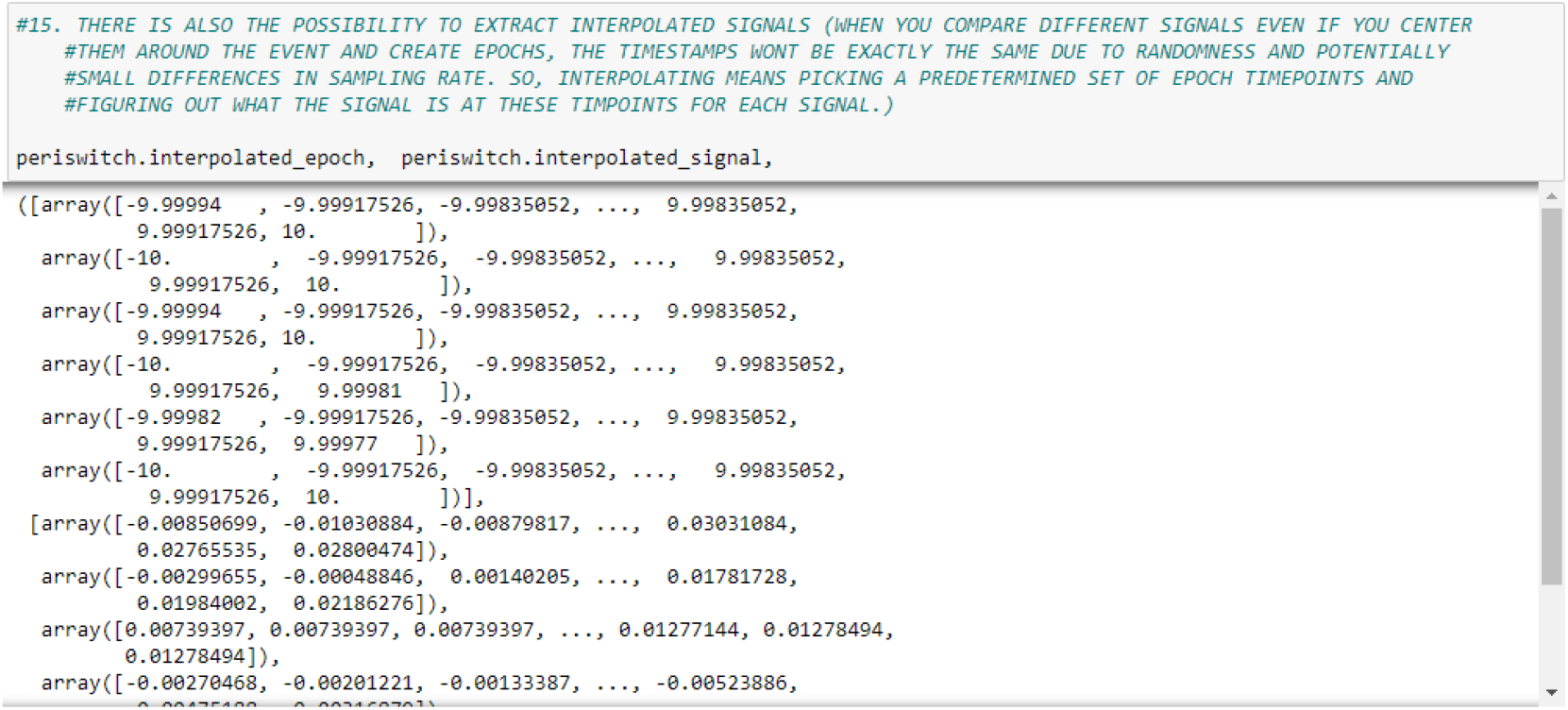

###### 4.1.2. Calcium signals by the Prelimbic cortex principal neurons in response to cocaine self-infusions

**Steps 1-2** are the same as 4.1.1.

**Step 3***(MultiSession Jupyter Notebook: #16)*: The Analyze module is called just as it was in the previous example, but for this specific case ‘inj1’ rather than ‘hled_on’ is used as the event, with a window of 3 seconds pre-event and 15 seconds post-event.

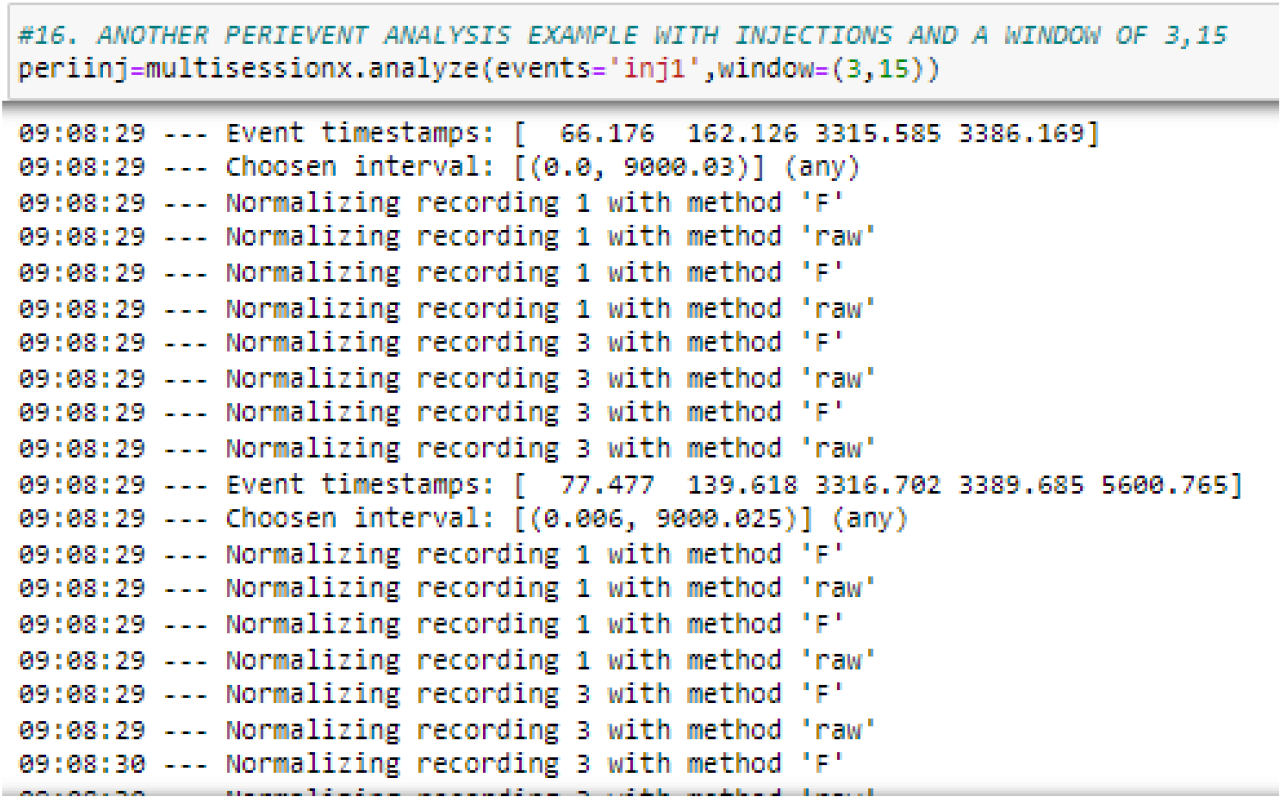

**Step 4***(MultiSession Jupyter Notebook: #17)*: Shown below is the plot of the perievent Z-scores. The signal, after the initial processing, is re-normalized using the average and standard deviation of the baseline period (the 3 seconds before the injection occurs).

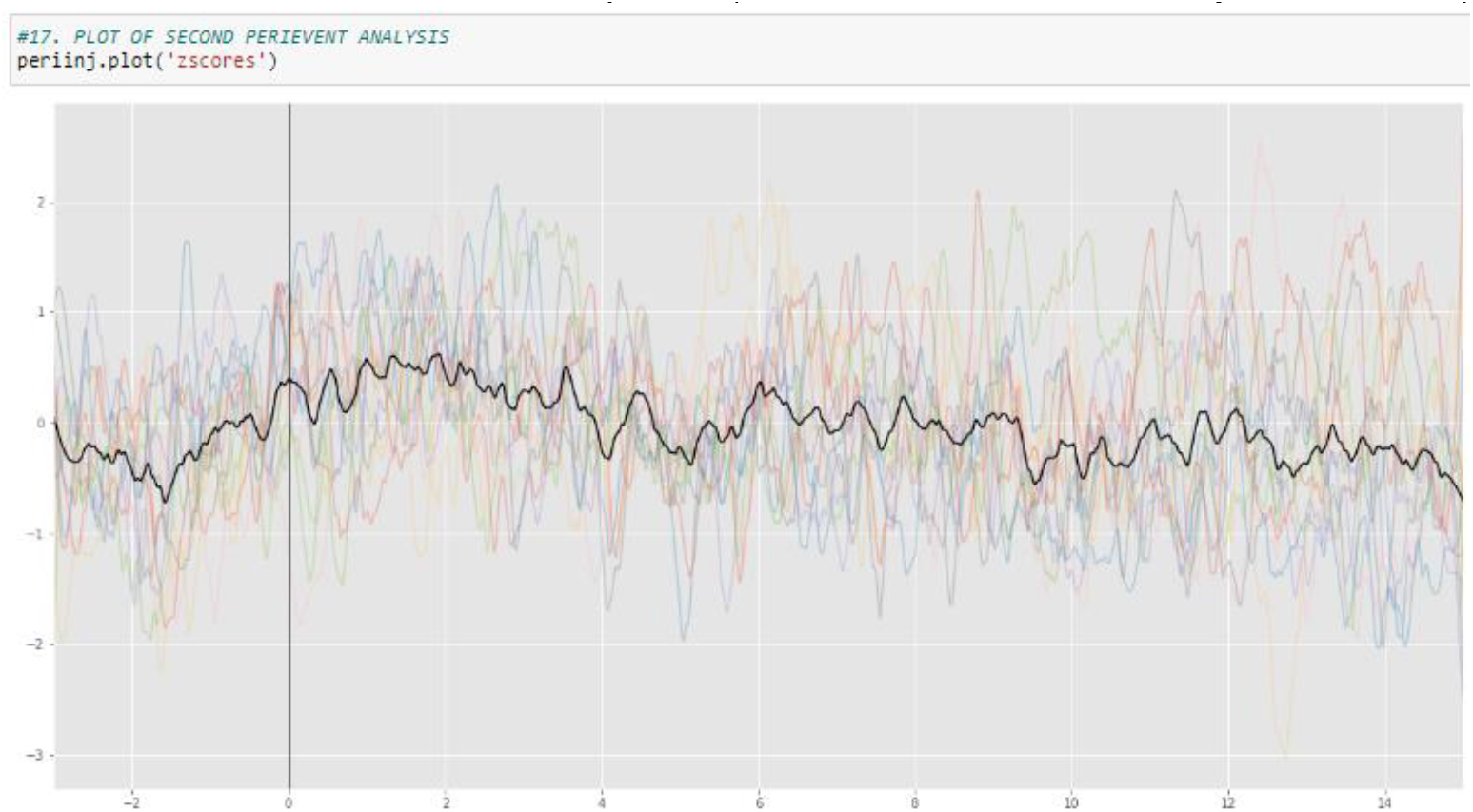

Further extraction of the perievent analysis can be done similar to what was shown in 4.1.1.

##### 4.2 Session

Below is a tutorial explaining how to use the Session module to analyze a single recording session.

As stated above, the Session module is used to analyze one behavioral and fiber data file in parallel. It uses the **behavior**, **fiber**, and **analyze** modules.

The corresponding Jupyter Notebook can be found and downloaded with the example file at: https://gitlab.com/inserm-u1215/pyfiber/-/tree/main/docs/notebooks/Session

###### 4.2.1. Analysis of fiber photometry signals in response to the first drug to no drug shift in a single self-administration session

**Step 1***(Session Jupyter Notebook: #1)*: Import *Pyfiber* in Jupyter Notebook.

**Step 2***(Session Jupyter Notebook: #2)*: Create a session object by calling pf.Session() and indicating the file path to the fiber and behavioral files in the parenthesis.

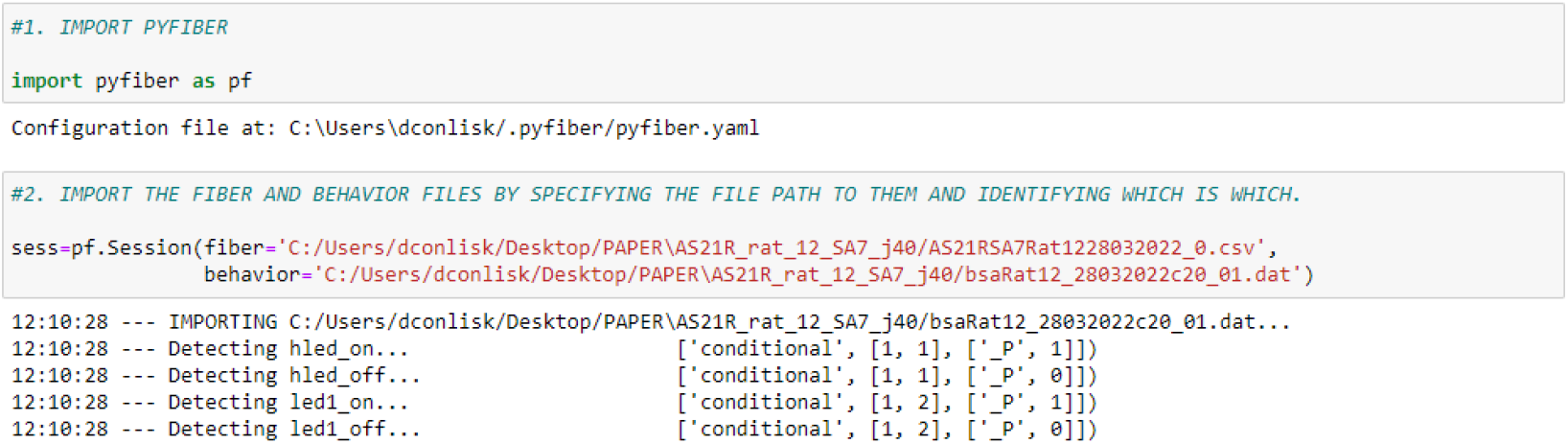

**Step 3***(Session Jupyter Notebook: #3)*: Identify timestamps of the events of interest. Shown here is the timestamps of injections in the first drug period and the timestamps of the switch from the first drug to the no drug period.

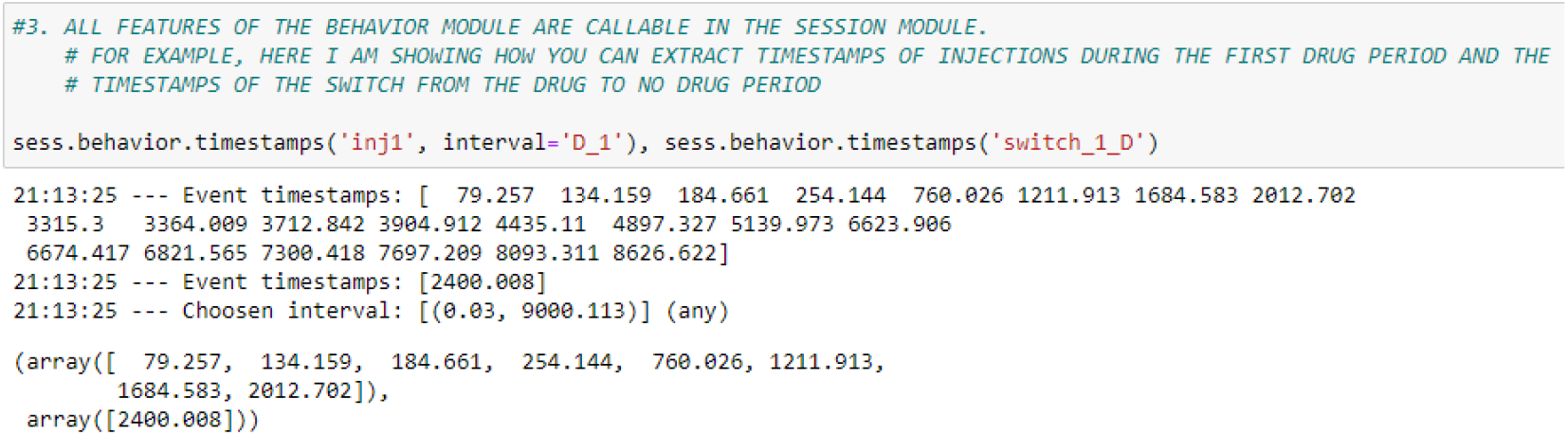

**Step 4***(Session Jupyter Notebook: #6)*: Using the timestamp of the switch, perievent analysis can be done by calling the Analyze module.

**Step 5***(Session Jupyter Notebook: #7)*: Like in the MultiSession module, the signal can be plotted by calling .plot().

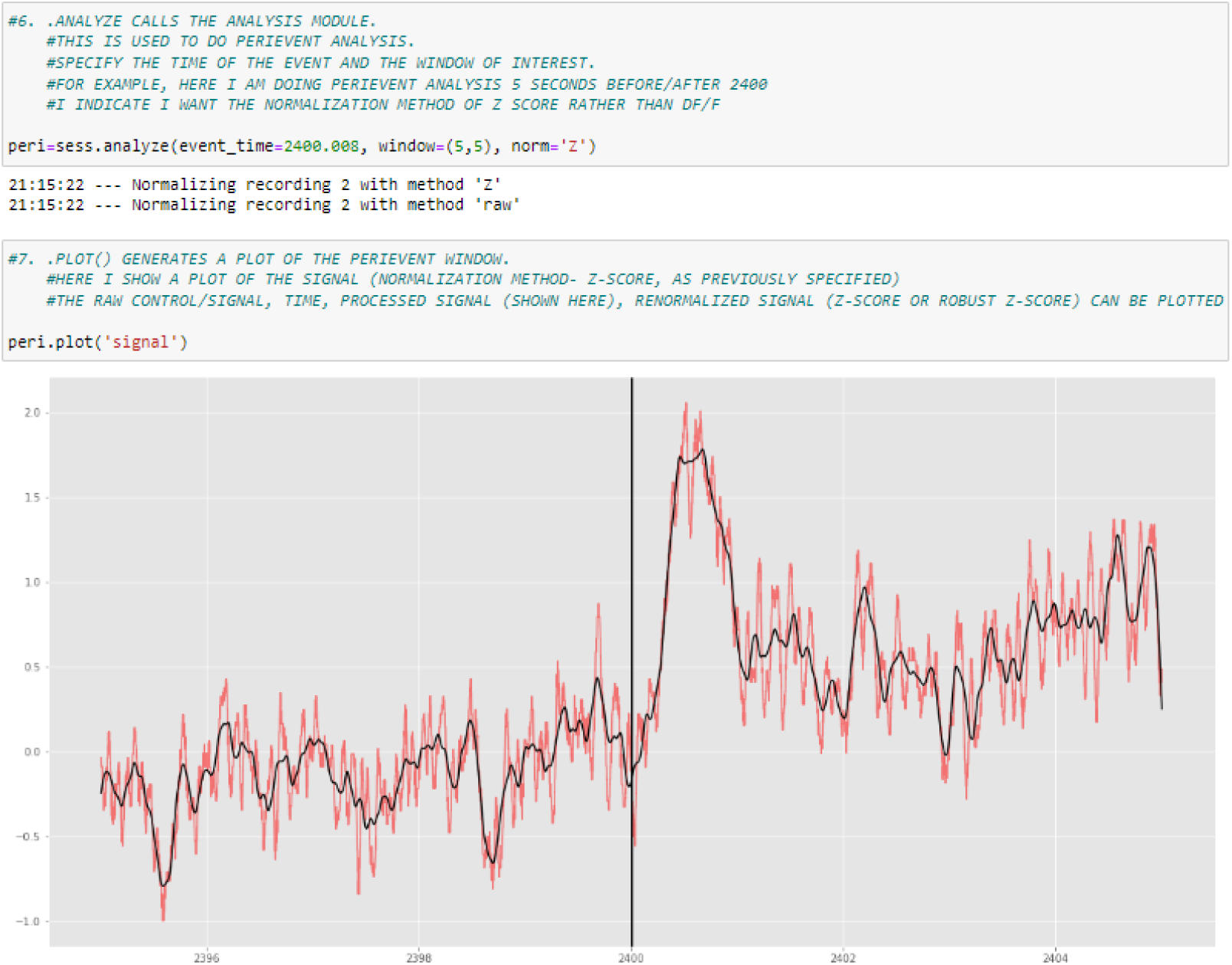

**Step 6***(Session Jupyter Notebook: #14)*: If the interest is comparing the peak frequency before and after the event occurs, this can be done by calling the analysis of interest within the perievent object.

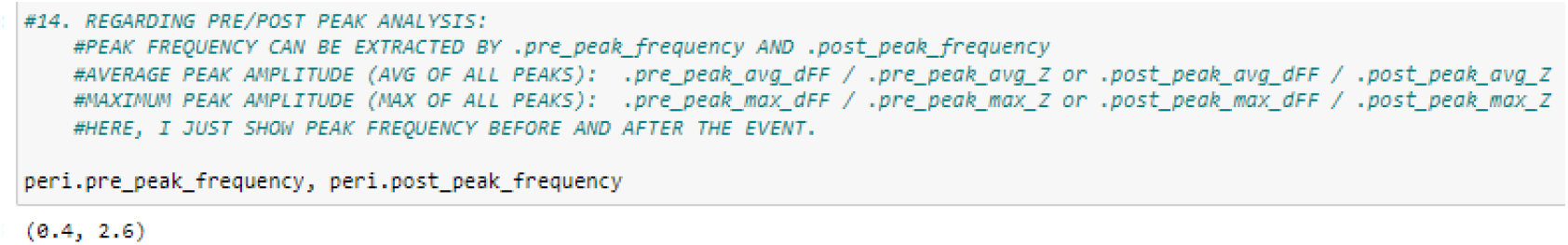

A table can be found in the SOM that contains a selection of additional features and commands associated with the Session module.

#### 5. Relevant information for all modules

##### 5.1 .help and .info

In all modules, there is guidance available by calling <obj>.help or <obj>.info.

##### 5.2 How to extract arrays

All of the data can be exported in multiple file formats depending on the user’s needs. This can be achieved by using Python’s built in functions or dedicated libraries such as pandas. An example below shows how to export the interpolated signal data for all events in a multiple analysis as a .csv file. The resulting .csv will contain a table with session names (by default folder names) as columns and timestamps as indices.

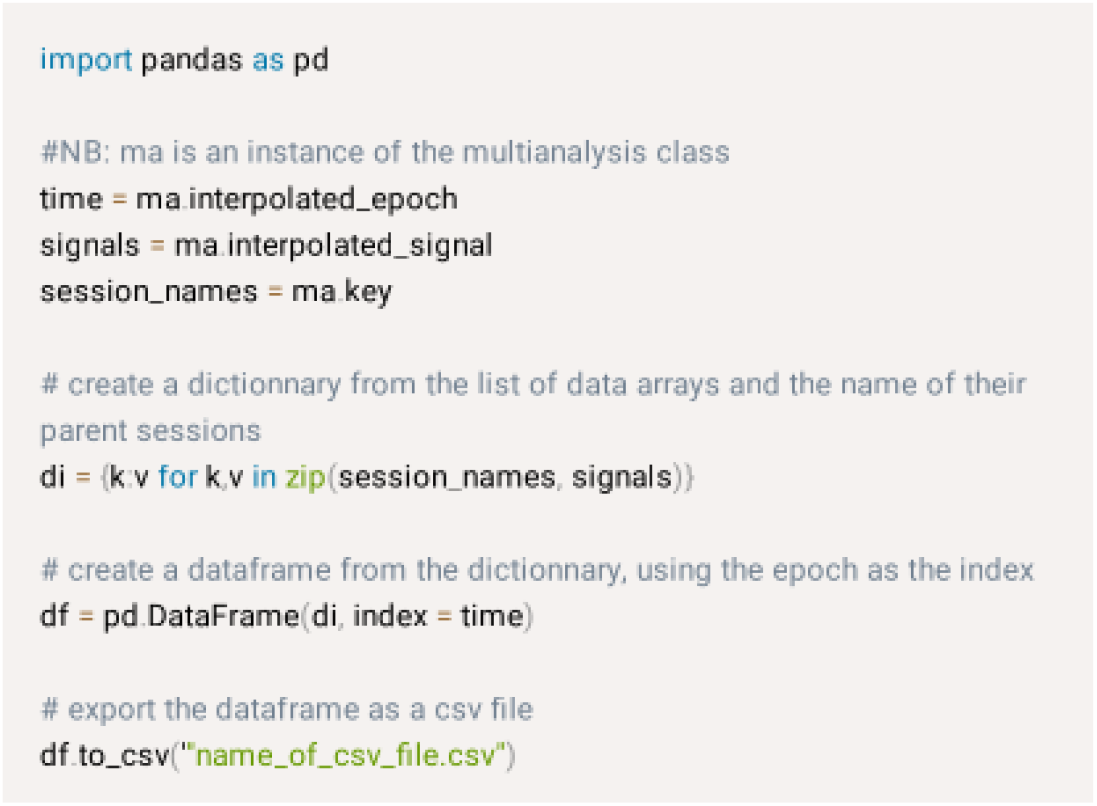

## DISCUSSION

### Benefits and constraints of fiber photometry

Many papers have now started to show the benefits of fiber photometry paired with certain aspects of operant behavior (Gioia & Woodward, 2021; Lafferty et al., 2020; Liu et al., 2020). The research pairing photometry with operant behavior is still in its infancy but has immense potential due to the multiple events occurring during operant behaviorx002D; but this also serves as a barrier that complicates data analysis. However, with appropriate analytic efficiency, the combination of techniques can uncover neural signatures surrounding drug/reward administration, lever pressing/nose-pokes, and conditioned/discriminative stimuli presentation/changes. However, this relies on proper identification and tagging of behavioral events, which can be complicated when using complex behavioral paradigms. This led us to create *Pyfiber*, a python library for joint behavioral and fiber photometry analysis with the idea of complex operant behavioral paradigms in mind (such as- but not limited to- intravenous self-administration).

### Current open-source tools for joint behavioral / fiber photometry analysis

While open source tools exist which aim to analyze fiber photometry recordings surrounding behavioral events, there were significant limitations which prevented us from using them. The pMAT program, despite having a very user-friendly interface, was quite rigid in regards to the types of analysis performed (Bruno et al., 2021). For example, signal processing was limited to ΔF/F, without the possibility of other techniques. Another limitation to pMAT is the fact that it was developed in Matlab, which requires a license. Even though the .exe is executable without Matlab, modifications of the pMAT program are not truly open source due to the requirement to have a Matlab license.

GuPPY, another open source fiber photometry tool, remedied a few of the aforementioned issues that were encountered with pMAT (Sherathiya et al., 2021). GuPPY was created in python, making it a true free and open source tool for fiber photometry analysis. Additionally, the tool implements many more options for analysis, such as transient frequency/amplitude analysis and other ways to process the raw fiber photometry signals. However, with GuPPY, the timestamps need to be extracted before analysis occurs, which can be a complicated step when using complex operant behavior paradigms.

### Justification of the creation of Pyfiber for joint behavioral / fiber photometry analysis

Various open source packages providing toolkits for fiber photometry analysis exist and the aforementioned programs have been made to provide easy-to-use softwares. However, specialized toolkits, often developed for task-specific applications often lack generalizability whereas more user-friendly projects appear to be more limited in their possibilities. As a growing field, fiber photometry still lacks a specialized code library that could provide the relevant tools for any type of fiber photometry data analysis projects. The modular approach of this project helps to achieve this aim and the object-oriented paradigm permits more flexibility when compared to a functional approach.

### Identification and categorization of behavioral events from raw ImetronicⓇ behavioral files

A necessary step to the combination of operant behavior and *in vivo* neuroimaging techniques is assuring that the events that are analyzed are categorically identical and can be pooled to uncover neurobiological signatures of the event of interest.

As there are many events occurring in self-administration including presentations of discriminative stimuli representing drug availability or unavailability, conditioned stimuli paired with reward delivery, reward delivery itself, drug seeking/taking behaviors, and more depending on the paradigm, a quick way to categorize events necessary for subsequent analysis is highly beneficial.

Identification of identical behavioral events can also be incredibly beneficial in the scope of other behavioral tests. For example, through the use of the Behavior module, quick identification of regressive and perseverative errors in operant cognitive flexibility tasks can be produced by quick modification and customization of the configuration file. By extension, the tool can be used to analyze data from 5-choice serial reaction time tasks (5-CSRTTs) to tag the behavioral events associated with impulsive action, attention, and reaction times.

Regardless of the behavioral paradigm, ensuring that the timestamps used for subsequent neurobiological analysis are identical is necessary to uncover neurobiological correlates of complex behaviors. The behavioral module within *Pyfiber* helps do that.

### Signal processing for fiber photometry analysis

Regarding the signal processing techniques, we implemented the most commonly used techniques- ΔF/F and Z-score. Introduction of a new type of signal processing, if necessary, can be done due to the flexibility of the library.

Another clear benefit of *Pyfiber* in the perspective of signal processing is the flexibility that is permitted with peak analysis. With *Pyfiber*, similar to GuPPY, the rolling window for transient detection can be modified. However, *Pyfiber*, unlike GuPPY, permits easier modifications to the thresholds necessary to define transients.

Signal analysis can then be compared between groups to identify differences in neural activity. The type of signal processing techniques that can be applied within *Pyfiber* was used to identify signatures of future stress susceptibility through the identification of differences in transient amplitude in medium spiny neurons in the nucleus accumbens (Muir et al., 2018). Similar studies can be done using *Pyfiber*’s fiber module to identify potential signatures of vulnerability to addiction, as well as other disorders.

### Selection of behavioral timestamps of interest and combination with photometry analysis

As previously discussed, the extraction and identification of identical behavioral events can be used to consequently analyze neural signatures surrounding these events. For example, timestamps of nose-pokes can be used as events of interest, as well as even choosing the specific operant response (nose-poke, lever press…) within the fixed ratio series by identification of the response in the series through modification of the configuration file. Being able to quickly identify each response in the series can be immensely beneficial in understanding neural signatures of drug/reward seeking. Notably, in previous data in the lab, it is uncovered that different nose-pokes in the FR5 series have different neural signatures (***data not shown***). However, this is not just applicable to our specific paradigm. In a progressive ratio test, the final nose-poke before the breakpoint was reached had the highest paranigral Ventral Tegmental Area response when compared to the other nose-pokes, uncovering subtle relationships between neuronal activity and behavioral responding in a motivational task (Parker et al., 2019).

Combining fiber photometry and operant behavior can also provide deeper insight on neural signatures surrounding reward delivery and aid in uncovering potential differences in neural signatures of reward-related mechanisms. In the scope of addiction, this can help identify potential differences in drug-induced sensitization. This can add to the current research that highlights differences in time-locked neural activity during drug consumption as a function of dependence during an operant self-administration task (Gioia & Woodward, 2021). Additionally, dissection of phasic and tonic neurotransmitter dynamics in response to drug delivery is also feasible through fiber photometry but has yet to be applied to paradigms with contingent drug delivery (Salinas et al., 2022).

Combining fiber photometry and operant behavior can also help uncover neural signatures in response to reward-related discriminative and conditioned stimuli. In the scope of drug-related behaviors, some studies have used conditioned place preference (CPP) to explore neural correlates of an exposure to a drug-associated context (Calipari et al., 2016; O’Neal et al., 2022). However, in CPP, animals do not freely self-administer the drug, which limits potential translatability.

Self-administration protocols are acknowledged to have higher translational value. Because animals self-administer the drug, the contextual stimuli paired with drug effects more similarly resemble drug-associated cues in human drug users. These conditioned stimuli, like drug associated cues in humans, strongly promote reinstatement of drug seeking behaviors (Shaham et al., 2003). Uncovering neural signatures in those who are exposed to these drug-paired cues and appropriately inhibit drug seeking compared to those who reinstate drug seeking behavior is yet to be performed. Identification that different experimental states (chronic stress or control) have different neural signatures in a cue-guided pavlovian autoshaping procedure substantiates the potential for different neural signatures of vulnerability to addiction to be uncovered in relation to conditioned drug stimuli encoding (Spring et al., 2021).

Dissection of neural correlates of other adaptive/maladaptive behaviors can be done through the combination of fiber photometry with operant tasks such as cognitive flexibility tasks or 5-CSRTT. The combination of complex cognitive testing and fiber photometry can contribute to the identification of the neural correlates involved in specific error types within cognitive flexibility paradigms as well as neural correlates related to impulsive action, attention, and reaction times through dissection of behavioral responses in 5-CSRTT trials (de Kloet et al., 2021).

Being able to pair fiber photometry with more complex decision-making processes will uncover important neural signatures in behaving animals with minimal manipulation of normal physiological states. These techniques can be especially useful when considering the pathological behavioral disruptions that occur in individuals with problematic substance use or other disorders that are currently modeled in rodents through complex behavioral paradigms.

We believe that *Pyfiber* can be a major asset to fluorescent imaging studies during operant behavior and will favor their still-limited implementation. This open source analytical python library helps eliminate some of the analytical difficulties that behavioral labs face when implementing bulk fluorescent imaging techniques, while still permitting the necessary flexibility that is required for a tool to be easily adapted to fit the needs of different laboratories.

### Limitations of Pyfiber

While *Pyfiber* can be useful to some, there are some limitations that need to be acknowledged and addressed in future updates. For example, a main limitation is that the Behavior module is only compatible with ImetronicⓇ software. As many researchers use behavioral apparati from other companies (notably, Med Associates), *Pyfiber* will be expanded to be able to convert raw data in other behavioral systems, however, the current version has not rectified this.

Likewise, currently *Pyfiber* only supports fiber photometry files from Doric systems. As fiber photometry data files are relatively similar across most systems, the conversion from other systems to the doric layout is quite simple. However, in a future version of *Pyfiber*, there will be the possibility to upload data types from other photometry systems.

Additionally, there is no graphical interface. This makes *Pyfiber* slightly more intimidating to use. However, providing Jupyter Notebooks that contain step-by-step instructions and example data on how to do the analysis alongside this paper and the documentation, users have adequate tools to be able to implement *Pyfiber* in their labs.

## AUTHOR CONTRIBUTIONS

DC and ED performed the behavioral experiments. MC developed *Pyfiber*. JFF, NW, and CH provided systems and experimental support. DC, MC and VDG wrote the manuscript. All authors reviewed the final version of the manuscript.

## FUNDING

This work was supported by l’Agence Nationale de la Recherche (ANR; grant ANR-10-EQX-008-1, EquipEx OptoPath to VD-G), by l’Institut National de la Santé et de la Recherche Médicale (INSERM) and the University of Bordeaux (UB). DC is a recipient of a PhD fellowship from the “Ecole Universitaire de Recherche” (EUR Neuro, Bordeaux Neurocampus). MC is a recipient of the 2021-2022 *Année Recherche* Program (DG-ARS, CHU Bordeaux).

We thank S. Laumond, J. Tessaire and the technical staff of the housing and experimental animal facility of the Neurocentre Magendie Inserm U1215. A special thanks to Cédric Dupuy for the remarkable care to our rats.

## REFERENCES

Bruno, C. A., O’Brien, C., Bryant, S., Mejaes, J. I., Estrin, D. J., Pizzano, C., & Barker, D. J. (2021). pMAT: An open-source software suite for the analysis of fiber photometry data. Pharmacology Biochemistry and Behavior, 201, 173093. https://doi.org/10.1016/j.pbb.2020.173093

Calipari, E. S., Bagot, R. C., Purushothaman, I., Davidson, T. J., Yorgason, J. T., Peña, C. J., Walker, D. M., Pirpinias, S. T., Guise, K. G., Ramakrishnan, C., Deisseroth, K., & Nestler, E. J. (2016). In vivo imaging identifies temporal signature of D1 and D2 medium spiny neurons in cocaine reward. Proceedings of the National Academy of Sciences, 113(10), 2726–2731. https://doi.org/10.1073/pnas.1521238113

de Kloet, S. F., Bruinsma, B., Terra, H., Heistek, T. S., Passchier, E. M. J., van den Berg, A. R., Luchicchi, A., Min, R., Pattij, T., & Mansvelder, H. D. (2021). Bi-directional regulation of cognitive control by distinct prefrontal cortical output neurons to thalamus and striatum. Nature Communications, 12(1), 1994. https://doi.org/10.1038/s41467-021-22260-7

Gioia, D. A., & Woodward, J. J. (2021). Altered Activity of Lateral Orbitofrontal Cortex Neurons in Mice following Chronic Intermittent Ethanol Exposure. Eneuro, 8(2), ENEURO.0503-20.2021. https://doi.org/10.1523/ENEURO.0503-20.2021

Holly, E. N., Davatolhagh, M. F., Choi, K., Alabi, O. O., Vargas Cifuentes, L., & Fuccillo, M. V. (2019). Striatal Low-Threshold Spiking Interneurons Regulate Goal-Directed Learning. Neuron, 103(1), 92–101.e6. https://doi.org/10.1016/j.neuron.2019.04.016

Lafferty, C. K., Yang, A. K., Mendoza, J. A., & Britt, J. P. (2020). Nucleus Accumbens Cell Type- and Input-Specific Suppression of Unproductive Reward Seeking. Cell Reports, 30(11), 3729–3742.e3. https://doi.org/10.1016/j.celrep.2020.02.095

Lerner, T. N., Shilyansky, C., Davidson, T. J., Evans, K. E., Beier, K. T., Zalocusky, K. A., Crow, A. K., Malenka, R. C., Luo, L., Tomer, R., & Deisseroth, K. (2015). Intact-Brain Analyses Reveal Distinct Information Carried by SNc Dopamine Subcircuits. Cell, 162(3), 635–647. https://doi.org/10.1016/j.cell.2015.07.014

Liu, Y., Jean-Richard-dit-Bressel, P., Yau, J. O.-Y., Willing, A., Prasad, A. A., Power, J. M., Killcross, S., Clifford, C. W. G., & McNally, G. P. (2020). The Mesolimbic Dopamine Activity Signatures of Relapse to Alcohol-Seeking. The Journal of Neuroscience, 40(33), 6409–6427. https://doi.org/10.1523/JNEUROSCI.0724-20.2020

Muir, J., Lorsch, Z. S., Ramakrishnan, C., Deisseroth, K., Nestler, E. J., Calipari, E. S., & Bagot, R. C. (2018). In Vivo Fiber Photometry Reveals Signature of Future Stress Susceptibility in Nucleus Accumbens. Neuropsychopharmacology, 43(2), 255–263. https://doi.org/10.1038/npp.2017.122

Nakai, J., Ohkura, M., & Imoto, K. (2001). A high signal-to-noise Ca2+ probe composed of a single green fluorescent protein. Nature Biotechnology, 19(2), 137–141. https://doi.org/10.1038/84397

O’Neal, T. J., Bernstein, M. X., MacDougall, D. J., & Ferguson, S. M. (2022). A Conditioned Place Preference for Heroin Is Signaled by Increased Dopamine and Direct Pathway Activity and Decreased Indirect Pathway Activity in the Nucleus Accumbens. The Journal of Neuroscience, 42(10), 2011–2024. https://doi.org/10.1523/JNEUROSCI.1451-21.2021

Parker, K. E., Pedersen, C. E., Gomez, A. M., Spangler, S. M., Walicki, M. C., Feng, S. Y., Stewart, S. L., Otis, J. M., Al-Hasani, R., McCall, J. G., Sakers, K., Bhatti, D. L., Copits, B. A., Gereau, R. W., Jhou, T., Kash, T. J., Dougherty, J. D., Stuber, G. D., & Bruchas, M. R. (2019). A Paranigral VTA Nociceptin Circuit that Constrains Motivation for Reward. Cell, 178(3), 653–671.e19. https://doi.org/10.1016/j.cell.2019.06.034

Salinas, A. G., Lee, J. O., Augustin, S. M., Zhang, S., Patriarchi, T., Tian, L., Morales, M., Mateo, Y., & Lovinger, D. M. (2022). Sub-second striatal dopamine dynamics assessed by simultaneous fast-scan cyclic voltammetry and fluorescence biosensor [Preprint]. Neuroscience. https://doi.org/10.1101/2022.01.09.475513

Shaham, Y., Shalev, U., Lu, L., de Wit, H., & Stewart, J. (2003). The reinstatement model of drug relapse: History, methodology and major findings. Psychopharmacology, 168(1–2), 3–20. https://doi.org/10.1007/s00213-002-1224-x

Sherathiya, V. N., Schaid, M. D., Seiler, J. L., Lopez, G. C., & Lerner, T. N. (2021). GuPPy, a Python toolbox for the analysis of fiber photometry data. Scientific Reports, 11(1), 24212. https://doi.org/10.1038/s41598-021-03626-9

Spring, M. G., Caccamise, A., Panther, E. A., Windsor, B. M., Soni, K. R., McReynolds, J. R., Wheeler, D. S., Mantsch, J. R., & Wheeler, R. A. (2021). Chronic Stress Prevents Cortico-Accumbens Cue Encoding and Alters Conditioned Approach. The Journal of Neuroscience, 41(11), 2428–2436. https://doi.org/10.1523/JNEUROSCI.1869-20.2021

Wang, W., Kim, C. K., & Ting, A. Y. (2019). Molecular tools for imaging and recording neuronal activity. Nature Chemical Biology, 15(2), 101–110. https://doi.org/10.1038/s41589-018-0207-0

Wang, Y., DeMarco, E. M., Witzel, L. S., & Keighron, J. D. (2021). A selected review of recent advances in the study of neuronal circuits using fiber photometry. Pharmacology Biochemistry and Behavior, 201, 173113. https://doi.org/10.1016/j.pbb.2021.173113

