## Supplemental Material for "*Pyfiber*: an open source python library that facilitates the merge of operant behavior and fiber photometry- focus on intravenous self-administration"

### SUPPLEMENTAL ONLINE MATERIALS

#### Supplemental Methods

##### Subjects

Male Sprague Dawley rats (CD, n=14) weighing 250-300g at their arrival were single housed in a temperature (22 ± 1°C) and humidity (60 ± 5%) controlled vivarium on a reverse-light cycle (ON 19h00, OFF 7h00). Food and water were provided *ad libitum*. Rats were handled daily before experimentation began.

##### Surgery

*Stereotaxic:* After 3-6 days of habituation, the animals were anesthetized with isoflurane gas (5% induction, 2% maintenance) and placed in a stereotaxic frame (Kopf). During surgery, body temperature was monitored and maintained at 37 degrees Celsius. Using a glass micropipette, 500 nL of GCaMP6f (Addgene, pENN.AAV.CamKII.GCaMP6f.WPRE.SV40- original titer: 2.30E+13, final titer: 2.30E+12) was injected into the prelimbic cortex (AP: +3.0, ML: +/- 0.6, DV: -3.5 from skull) at a rate of approximately 250 nanoliters/minute. The micropipette was allowed to stay in place for 10 minutes to allow for diffusion. A hollow blunted tip 27G needle (outer diameter 410 micrometers) was lowered to the placement of the fiber then removed to prevent tissue accumulation under the recording area of the fiber. Lastly, the fiber (400 µm core, 0.48 NA, 5 mm in length; Doric Lenses) was inserted and secured with Metabond (C&B Metabond, Parkell). The rats were treated post-op with the analgesic metacam (1 mg/kg, SC).

*Intravenous catheterization:* Four weeks after the stereotaxic surgery, the rats underwent intravenous catheterization. A silastic catheter (internal diameter = 0.28 mm; external diameter = 0.61 mm; dead volume = 12 µl) was implanted in the right jugular vein under isoflurane anesthesia. The proximal end of the catheter was inserted into the right atrium, passed under the skin, and the base emerged from the mid scapular region. Rats were treated post-op with metacam (1 mg/kg, SC) and allowed to recover for 5 to 9 days after surgery before self-administration began.

##### Histology

At the end of the experiment, the rats were deeply anesthetized with an intraperitoneal injection of a mix of pentobarbital and lidocaine (200 mg/kg pentobarbital / 20 mg/kg lidocaine). The animals were transcardially perfused with 4% paraformaldehyde (PFA) and then their brains were removed and stored in 4% PFA until they were sliced. The brains were sliced (slice thickness 50 μm) and immediately placed on glass slides. Proper fiber placement and adequate virus expression was analyzed using a Nikon Eclipse inverted microscope (Ti-U) **(Figure S2)**.

**Intravenous self-administration training**

*Basal self-administration session*. All experiments were performed in the dark phase of the light/dark cycle. The daily sessions consisted of three drug periods (40 min) separated by two no drug periods (15 min).  Drug periods were characterized by illumination of the chamber by a blue light (LED2) while no drug periods were characterized by illumination of the chamber by a white house light (HLED) **(Figure 1A)**. Inactive nosepokes during the duration of the sessions were recorded but had no scheduled consequences and active nosepokes during the no drug period were recorded but had no scheduled consequences. Animals were trained on a fixed ratio 3 (FR3) of responding for the first three days of self-administration (max infusions 25, 25, 30, respectively), then continued on an FR5 ratio of responding for the duration of the experiment (max infusions 30, then 35). Once the animals reached the necessary number of nosepokes for infusion, the white cue light (LED1) illuminated for four seconds, with the pump being activated (delivering 46 microliters, 0.8 mg/kg/inf) one second after the illumination of the white light. Then, both the white cue light and blue light were turned off during the 40 second timeout period **(Figure 1B)**. Active and inactive responding during the timeout period were recorded but had no scheduled consequences.

**Supplemental Figures**

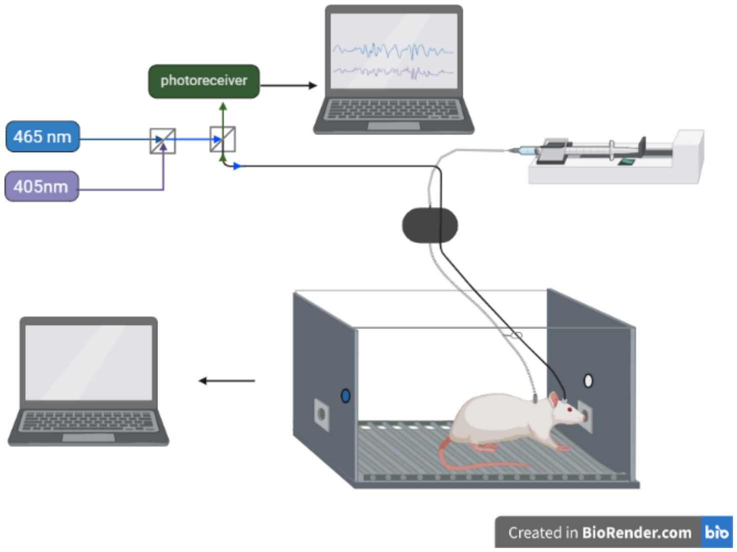


**Figure S1:** Schematic of the fiber photometry and behavioral systems. Fiber photometry data is collected by the Doric Lenses software system while the operant behavior data is collected by the Imetronic® system. The fiber photometry system initiates and terminates recordings based on TTL pulses generated from the Imetronic® system. Created by BioRender.com.

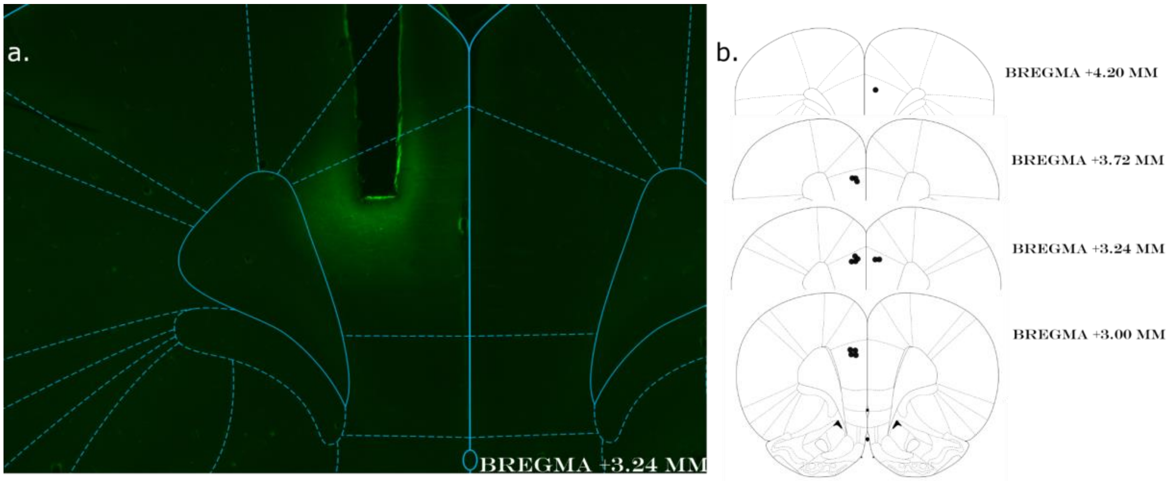


**<u>Figure S2:</u> A.** Example of expression of the GCaMP6f in the PL attested by the green fluorescence emitted by the circularly permuted green fluorescent protein (cpGFP), and location of the tip of the optical fiber. B. Placements of the optical fiber in the 14 tested rats targeting the PL from AP +4.2 to AP +3 mm.

### Supplementary information on *Pyfiber* modules

#### 1. Application of the Fiber and Behavior (MultiBehavior and Behavior) modules

##### 1.1. Fiber Module

An example Jupyter Notebook that can be used as a reference for the benefits of the Fiber module can be found and downloaded with the example file at:

**https://gitlab.com/inserm-u1215/pyfiber/-/tree/main/docs/notebooks/Fiber**

##### 1.2. Behavior Module

###### 1.2.1. Behavior

An example Jupyter Notebook that can be used as a reference for the benefits of the behavior module can be found and downloaded with the example file at:

**https://gitlab.com/inserm-u1215/pyfiber/-/tree/main/docs/notebooks/Behavior**

###### 1.2.2. MultiBehavior

An example Jupyter Notebook that can be used as a reference for the benefits of the MultiBehavior module can be found and downloaded with the example files at:

**https://gitlab.com/inserm-u1215/pyfiber/-/tree/main/docs/notebooks/MultiBehavior**

### 2. Features and Commands associated with the different modules

#### 2.1. MultiSession Module

Below is a table containing a selection of the most pertinent features and the commands associated with the MultiSession module:

multisessionx = pyfiber.MultiSession('path_to_the_folder_with_subfolders_containing_a_dat_and_csv_file')

| Method | Feature | Command example | Jupyter Notebook Title |
|---|---|---|---|
| Details | Shows a list containing the experiment, rat number, experiment type, session number, ID, and session tag | multisessionx.details | Multisession #3 |
| Folder | Shows the location of the folder that contains the subfolders containing the data files | multisessionx.folder | Multisession #4 |
| Names | Shows the session tags | multisessionx.names | Multisession #5 |
| Removed | Shows removed sessions (due to us having a problem with the duration of the Imetronic system- theoretically no one else will have this problem) | multisessionx.removed | |
| Sessions | A dictionary containing the session tag and the locations for the behavior and fiber data files | multisessionx.sessions | Multisession #6 |
| Compare Behavior | Output is a heat map and a plot containing a visual representation of the differences in behavior (a single event). Used to visualize behavioral differences between sessions/rats. | multisessionx.compare_behavior('np1') | Multisession #7 |
| **Analyze** | | | |
| Analyze | Prepare analyzed fiber data related to defined perievent window. The analysis needs to be put in a variable which is used to get graphs or dataframes. | MultiAnalyze = multisession.analyze('switch_1_D', window=(10,10)) | Multisession #8, #16 |
| Plot | Plot the perievent analysis (either time, raw control, raw signal, processed signal, z-score, or robust z-score). | MultiAnalyze.plot('time')<br>MultiAnalyze.plot('raw_control')<br>MultiAnalyze.plot('raw_signal')<br>MultiAnalyze.plot('signal')<br>MultiAnalyze.plot('zscores')<br>MultiAnalyze.plot('rob_zscores') | Multisession #9, #17 |

| **Data** | **Return a dataframe with each event as a row and the columns containing all information below in red, which can also be called individually** | **MultiAnalyze.data** | **Multisession #10, #18** |
|---|---|---|---|
| Verification of filepath | The filepath for the behavioral or fiber data files<br>Can also be called without .data | MultiAnalyze.fiberfile<br>MultiAnalyze.behaviorfile | Multisession #11 |
| Normalization method | Verification of the normalization method used (should be what is indicated in the configuration file unless otherwise indicated | MultiAnalyze.normalisation | Multisession #11 |
| Verification of event time | Output is the time of the event | MultiAnalyze.event_time | Multisession #11 |
| Verification of perievent window | Output is the perievent window | MultiAnalyze.window | Multisession #11 |
| Sampling rate of recordings used | Output is the sample rate of the recording used | MultiAnalyze.sampling_rate | Multisession #11 |
| Recording number of recording used | Output is the recording number of the recording used for the perievent analysis | MultiAnalyze.data.rec_number | Multisession #11 |
| Pre/post peak frequency | Output is the frequency of peaks/second either pre or post event | MultiAnalyze.pre_peak_frequency<br>MultiAnalyze.data.post_peak_frequency | Multisession #11 |
| Average pre/post peak amplitude | Output is the average amplitude of the peaks either pre or post event (either ΔF/F or Z-score) | MultiAnalyze.pre_peak_AVG_dFF<br>MultiAnalyze.post_peak_AVG_dFF<br>MultiAnalyze.pre_peak_AVG_Z<br>MultiAnalyze.post_peak_AVG_Z | Multisession #11 |
| Maximum pre/post peak amplitude | Output is the maximum peak amplitude either pre or post event (either ΔF/F or Z-score) | MultiAnalyze.pre_peak_max_ dFF<br>MultiAnalyze.post_peak_max_dFF<br>MultiAnalyze.pre_peak_max_Z<br>MultiAnalyze.post_peak_max_Z | Multisession #11 |
| Pre/post AUC | Output of the area under the curve of the pre/post signal of choice (raw (calcium dependent channel), processed signal, robust Z-score, or Z-score) | MultiAnalyze.pre_raw_AUC<br>MultiAnalyze.post_raw_AUC<br>MultiAnalyze.preAUC<br>MultiAnalyze.postAUC<br>MultiAnalyze.preRZ_AUC<br>MultiAnalyze.postRZ_AUC<br>MultiAnalyze.preZ_AUC<br>MultiAnalyze.postZ_AUC | Multisession #11 |

| | | | |
|---|---|---|---|
| Pre/post average values | Output is the average value of the pre/post signal of choice (raw(preprocessed), robust Z-score, or Z-score) | MultiAnalyze.preAVG_dF<br>MultiAnalyze.postAVG_dF<br>MultiAnalyze.preAVG_RZ<br>MultiAnalyze.postAVG_RZ<br>MultiAnalyze.preAVG_Z<br>MultiAnalyze.postAVG_Z | Multisession #11 |
| Full Data | Return a dataframe with each event as a row and the columns containing all information above in red and the dataframes below in green (which can also be called individually by calling MultiAnalyze.full_data._____) | MultiAnalyze.full_data | Multisession #12 |
| Pre/post event dataframes | Output of a dataframe containing the time, raw (control or signal), processed data, or re-normalized data (Robust Z-scores or Z-scores) selectively either before or after the event occurs | MultiAnalyze.pre_time<br>MultiAnalyze.post_time<br>MultiAnalyze.pre_raw_ctrl<br>MultiAnalyze.post_ raw_ctrl<br>MultiAnalyze.pre_raw_sig<br>MultiAnalyze.post_ raw_sig<br>MultiAnalyze.preevent<br>MultiAnalyze.postevent<br>MultiAnalyze.pre_Rzscores<br>MultiAnalyze.post_Rzscores<br>MultiAnalyze.pre_zscores<br>MultiAnalyze.post_zscores | Multisession #13 |
| Raw Data | An array containing the time and raw data (raw signal and control channels) or the time and processed signal for the entire recording that was used for the perievent analysis | MultiAnalyze.rawdata<br>MultiAnalyze.recordingdata | Multisession #14 |
| Export individual signals | An array containing only the time, raw control channel, calcium dependent channel, processed data, or re-normalized data (Robust Z-scores or Z-scores) selectively during the perievent time period | MultiAnalyze.time<br>MultiAnalyze.raw_control<br>MultiAnalyze.raw_signal<br>MultiAnalyze.signal<br>MultiAnalyze.rob_zscores<br>MultiAnalyze.zscores | Multisession #13 |
| Data | Perievent data (time and processed signal (ΔF/F or Z-score) | MultiAnalyze.full_data.data | Multisession #13 |
| Export average signals | An array containing the mean values of all recordings | MultiAnalyze.EPOCH<br>MultiAnalyze.RAW_CONTROL<br>MultiAnalyze.RAW_SIGNAL<br>MultiAnalyze.ROB_ZSCORES<br>MultiAnalyze.SIGNAL<br>MultiAnalyze.TIME<br>MultiAnalyze.WINDOW<br>MultiAnalyze.ZSCORES | |

| | | | |
|---|---|---|---|
| Export interpolated signals | An array with interpolated signals (due to slight differences that exist between different recordings) | MultiAnalyze.interpolated_epoch<br>MultiAnalyze.interpolated_raw_control<br>MultiAnalyze.interpolated_raw_signal<br>MultiAnalyze.interpolated_rob_zscores<br>MultiAnalyze.interpolated_signal<br>MultiAnalyze.interpolated_time<br>MultiAnalyze.interpolated_zscores | Multisession #15 |

#### 2.2. Session Module

Below is a table containing a selection of the most pertinent features and the commands associated with the Session module:

session = pf.Session('path_to_the_folder_with_dat_and_csv_file')

| Method | Feature | Command example | Jupyter Notebook Title |
|---|---|---|---|
| | | **Behavior** | |
| All Behavior Methods (see table 1) | All features available in pyfiber.Behavior | session.behavior.______ | Session #3, #4 |
| | | **Fiber** | |
| All Fiber Methods (see table 3) | All features available in pyfiber.Fiber | session.fiber.______ | Session #5 |
| | | **Analyze** | |
| Analyze | Prepare analyzed fiber photometry data related to defined event extracted from the perievent window. Need to put it in a variable and then use it to get graphs or dataframes. | objAnalyze = session.analyze('np1', window=(0.5, 0.5)) | Session #6 |
| Plot | Plot the perievent analysis (either time, raw control, raw signal, processed signal, z-score, or robust z-score) | objAnalyze.plot('time')<br>objAnalyze.plot('raw_control')<br>objAnalyze.plot('raw_signal')<br>objAnalyze.plot('signal')<br>objAnalyze.plot('zscores')<br>objAnalyze.plot('rob_zscores') | Session #7 |
| Smooth | Outputs a smoothed array (using the Savitzky-Golay filter) | objAnalyze.smooth('time')<br>objAnalyze. smooth ('raw_control')<br>objAnalyze. smooth ('raw_signal')<br>objAnalyze. smooth ('signal')<br>objAnalyze. smooth ('zscores')<br>objAnalyze. smooth ('rob_zscores') | Session #8 |
| Data | Perievent data (time and processed signal (dF/F or Z-score) | objAnalyze.data | Session #9 |
| Full Data Frame- Raw and/or processed | An array containing the time and raw data (raw signal and control channels) or the time and processed signal for the entire recording that was used for the perievent analysis | objAnalyze.rawdata<br>objAnalyze.recordingdata | Session #10 |

| | | | |
|---|---|---|---|
| Export signal | An array containing only the time, raw control channel, calcium dependent channel, processed data, or re-normalized data (Robust Z-scores or Z-scores) selectively during the perievent time period | objAnalyze.time<br>objAnalyze.raw_control<br>objAnalyze.raw_signal<br>objAnalyze.signal<br>objAnalyze.rob_zscores<br>objAnalyze.zscores | Session #11 |
| Pre/post event dataframes | Output of a dataframe containing the time, raw (control or signal), processed data, or re-normalized data (Robust Z-scores or Z-scores) selectively either before or after the event occurs | objAnalyze.pre_time<br>objAnalyze.post_time<br>objAnalyze.pre_raw_ctrl<br>objAnalyze.post_ raw_ctrl<br>objAnalyze.pre_raw_sig<br>objAnalyze.post_ raw_sig<br>objAnalyze.preevent<br>objAnalyze.postevent<br>objAnalyze.pre_Rzscores<br>objAnalyze.post_Rzscores<br>objAnalyze.pre_zscores<br>objAnalyze.post_zscores | Session #12 |
| Pre/post AUC | Output of the area under the curve of the pre/post signal of choice (raw (calcium dependent channel), processed signal, robust Z-score, or Z-score) | objAnalyze.pre_raw_AUC<br>objAnalyze.post_raw_AUC<br>objAnalyze.preAUC<br>objAnalyze.postAUC<br>objAnalyze.preRZ_AUC<br>objAnalyze.postRZ_AUC<br>objAnalyze.preZ_AUC<br>objAnalyze.postZ_AUC | Session #13 |
| Pre/post average values | Output is the average value of the pre/post signal of choice (raw(preprocessed), robust Z-score, or Z-score) | objAnalyze.preAVG_dF<br>objAnalyze.postAVG_dF<br>objAnalyze.preAVG_RZ<br>objAnalyze.postAVG_RZ<br>objAnalyze.preAVG_Z<br>objAnalyze.postAVG_Z | Session #13 |
| Pre/post peak frequency | Output is the frequency of peaks/second either pre or post event | objAnalyze.pre_peak_frequency<br>objAnalyze.post_peak_frequency | Session #14 |
| Average pre/post peak amplitude | Output is the average amplitude of the peaks either pre or post event (either ΔF/F or Z-score) | objAnalyze.pre_peak_avg_dFF<br>objAnalyze.post_peak_avg_dFF<br>objAnalyze.pre_peak_avg_Z<br>objAnalyze.post_peak_avg_Z | Session #14 |
| Maximum pre/post peak amplitude | Output is the maximum peak amplitude either pre or post event (either ΔF/F or Z-score) | objAnalyze.pre_peak_max_dFF<br>objAnalyze.post_peak_max_dFF<br>objAnalyze.pre_peak_max_Z<br>objAnalyze.post_peak_max_Z | Session #14 |
| Verification of filepath | The filepath for the behavioral or fiber data files | objAnalyze.fiberfile<br>objAnalyze.behaviorfile | Session #15 |
| Normalization method | Verification of the normalization method used (should be what is indicated in the configuration file unless otherwise indicated | objAnalyze.normalisation | Session #16 |
| Verification of event time | Output is the time of the event | objAnalyze.event_time | Session #17 |
| Verification of perievent window | Output is the perievent window | objAnalyze.window | Session #17 |

| | | | |
|---|---|---|---|
| Sampling rate of recordings used | Output is the sample rate of the recording used | objAnalyze.sampling_rate | Session #18 |
| Recording number of recording used | Output is the recording number of the recording used for the perievent analysis | objAnalyze.rec_number | Session #18 |

#### 2.3. Fiber Module

Below is a table containing a selection of the most pertinent features and the commands associated with the Fiber module:

fiber = pf.Fiber('path_to_the_csv_file')

| Method | Feature | Command example | Jupyter Notebook Title |
|---|---|---|---|
| Plot | Show graphs with Fiber data. | fiber.plot() | Fiber #3 |
| Plot transients | Show a graph with transients among time. | fiber.plot_transients() | Fiber #4 |
| Export to CSV | Export data to a CSV file. This reformats the fiber data to the configuration that is used in GuPPY. | fiber.to_csv() | Fiber #5 |
| Normalize | Normalize data with specified method. Can be with either ΔF/F ('F') or Z-scores ('Z'). Can be specific recordings, but the default is all. | fiber.norm(method='F', rec=1) | Fiber #6, #7 |
| Peaks | Return a dictionary containing a data frame with peak data (timestamps and amplitude (ΔF/F and Z-score)) for all recordings. | fiber.peaks | Fiber #8 |
| PeakFA | After indicating the boundaries, it returns the peak frequency, average, and maximum amplitude of peaks (both ΔF/F and Z-score) | fiber.peakFA(2400, 2405) | Fiber #9 |
| Get | Extracts data array for a specific column of a recording. Can be either 'time,' 'signal,' or 'control.' | fiber.get('signal') | Fiber #10 |

**2.4. Behavior Module**

2.4.1. Behavior

Below is a table containing a selection of the most pertinent features and the commands associated with the Behavior module:

behavior = pf.Behavior('path_to_the_.dat_file')

| Method | Feature | Command example | Jupyter Notebook Title |
|---|---|---|---|
| Summary | Show a graphical overview of main events and intervals (as defined in the configuration file). | behavior.summary() | Behavior #5 |
| Figure | Plot event of interest along time. | behavior.figure('inj1') | Behavior #6 |
| Total | View count of all events | behavior.total | Behavior #7 |
| Raw | View raw data | behavior.raw | Behavior #8 |
| Data | Output is an array containing all events and when they occur (in which intervals) | behavior.data | Behavior #9 |
| Timestamps | Return array with timestamps of an event according to filters. | behavior.timestamps('inj1', interval = ('D_1')) | Behavior #10, #11 |
| Export Timestamps | Same as Timestamps with a graphical representation of events and inclusion/exclusion criteria. | behavior.export_timestamps('inj1', interval = ('D_1')) | Behavior #12 |
| Events | If nothing is in the (), there will be an output containing all the events of the session.<br>If an event is specified (e.g, 'inj1'), the output will be all events that occur during fiber recording (when TTL1=ON). | behavior.event()<br><br>behavior.event('inj1') | Behavior #13 |
| Intervals | Return list of all intervals defined in the configuration file. | behavior.intervals() | Behavior #14 |
| Specific Data visualization | Return a dataframe summarizing the Imetronic data from the dat file for a specific event. | behavior.get('INJ1') | Behavior #15 |
| Movement | Produces a heatmap of localization in the box (in our case, left or right as we only have two beams). | behavior.movement() | Behavior #16 |

2.4.2. MultiBehavior

Below is a table containing a selection of the most pertinent features and the commands associated with the MultiBehavior module:

multibehavior = pf.MultiBehavior('path_to_the_repertory_with_dat_files')

| Method | Feature | Command example | Jupyter Notebook Title |
|---|---|---|---|
| Count | Returns a dataframe with the cumulative number of events per second. | multibehavior.count('np1') | MultiBehavior #3 |
| Cumul | Show a graph with events accumulated over time animal per animal. | multibehavior.cumul('np1') | MultiBehavior #4 |
| Show Rate | Show rate for all sessions for any given event (default window is 120s). Also, shows a graph including the $15^{th}$, $50^{th}$, and $85^{th}$ percentiles. | multibehavior.show_rate('np1') | MultiBehavior #5 |
| Summary | Same as summary of Behavior module but for all the analyzed sessions. | multibehavior.summary() | MultiBehavior #6 |
| Timestamps of events | Shows a dataframe with each row as a .dat file and the columns containing the timestamps of the event or interval of interest. | multibehavior.inj1<br>multibehavior.led1_on<br>(and all other events or intervals within the configuration file) | MultiBehavior #7 |
